# Capturing Regional Variation in Aortic Mechanics: Dual-Estimation Method for Material Parameter Identification and Biological Correlation

**DOI:** 10.64898/2026.05.29.728673

**Authors:** Ricardo Doll Lahuerta, Ayumi A. Miyakawa, Marina J. S. Maizato, Renato Crajoinas, Bruno Durante da Silva, José Eduardo Krieger, Eduardo Moacyr Krieger, Idágene A. Cestari

## Abstract

The aorta shows significant regional variation in geometry and composition. This complexity makes numerical modeling challenging, as it requires identifying material parameters. Typically, the Holzapfel-Gasser-Ogden model is used. However, it suffers from nonuniqueness and sensitivity to outliers, which can obscure biological variation. In addition, standard compressible formulations with a volumetric-isochoric split fail to couple volumetric and anisotropic responses. To address these issues, a regularized dual-estimation framework was introduced. This framework combines a global baseline estimator with local refinement while maintaining structural material continuity. Furthermore, it uses a Modified Anisotropic model to improve the representation of compressibility physics. For validation, the approach included uniaxial extension and protein quantification from Wistar rats. The results show that the proximal ascending/aortic-arch segment is most compliant at low stretch, whereas the abdominal aorta stiffens earlier and becomes fiber-dominated at lower stretch levels. Notably, these trends align directionally with regional composition. However, the fitted stress components are model-based descriptors rather than direct measurements of individual constituents.

## 1 Introduction

Knowledge of the material properties of the aortic wall may contribute to the design of new drugs and treatments for cardiovascular diseases, as these properties offer important insights into the complex hemodynamics and biomechanics of the aorta. The structural behavior of the aortic wall is governed by complex interactions between its microstructural components, which exhibit regional differences along its length. The stiffness and composition of fibrous proteins, for instance, can vary substantially between the proximal and distal aortic segments. Such variations have physiological implications for the progression of vascular diseases and the mechanical response of the aorta under loading conditions. [31].

### 1.1 Aortic Wall Structure and Function

The aortic wall is a sophisticated trilaminate structure, with each layer possessing distinct structural attributes essential for maintaining overall aortic function [31]. The tunica intima, the innermost layer, comprises endothelial cells overlying a subendothelial matrix rich in elastin and collagen, which facilitates smooth blood flow and participates in regulating vascular tone. The tunica media, a thick middle layer, is predominantly composed of elastin and circumferentially arranged layers of spindle-shaped smooth muscle cells (SMCs). These SMCs constitute approximately 40 − 70% of the wall’s thickness and, in conjunction with collagen fibers, endow the aorta with its characteristic elasticity and tensile strength. Finally, the tunica adventitia, the outermost layer, consists of collagen-rich connective tissue and fibroblasts, featuring thick, wavy bundles of collagen fibers that prevent overstretching and potential rupture under high blood pressure.

A key aspect is the significant variation in the microstructure along the longitudinal axis of the aorta. For instance, in the ascending aorta, elastin fibers are arranged in a nearly isotropic pattern, whereas collagen fibers tend to be aligned more circumferentially. This specific architecture enables the proximal aorta to effectively manage the heart’s pulsatile flow, storing systolic volume as potential energy and releasing it during diastole to maintain a relatively continuous blood flow to the periphery. This elastic dampening, known as the Windkessel effect, was first observed by Hales in 1733 [61]. In contrast, the abdominal aorta has a more anisotropic microstructure, with collagen fibers typically arranged in a helical pattern, which increases the resistance to both circumferential and axial deformations, thereby enhancing local stiffness and reducing elasticity. The SMCs in the abdominal aorta are also relatively more abundant and play vital roles in regulating regional blood flow and maintaining systemic blood pressure. A thorough understanding of these structural gradients is indispensable for accurate biomechanical modeling [32].

### 1.2 Existing Models and Limitations

The biomechanics of the aortic wall have been investigated through a combination of theoretical, computational, and experimental approaches, combining numerical modeling with empirical data [16, 41, 28, 63, 27, 26, 20, 54, 64, 10]. A widely adopted constitutive relation for describing the nonlinear anisotropic behavior of soft biological tissues is the hyperelastic model proposed by Holzapfel et al. [26], often referred to as the Holzapfel–Gasser–Ogden (HGO) model. This model is based on a strain-energy function that is partitioned into an isotropic component, which represents the ground matrix, and an anisotropic component, which accounts for the behavior of embedded fibers. The isotropic part typically captures the response of the amorphous matrix material, while the anisotropic part models the directional reinforcement provided by collagen fibers, together describing the tissues stress-strain response.

Historically, arterial tissue has been modeled as an incompressible material, a simplification supported by early findings such as those by Carew et al. [9], who reported negligible volume changes (*<* 0.06%) under physiological pressures. However, this hypothesis has been revisited due to recent experimental evidence indicating that the arterial wall can exhibit volumetric compressibility, with volume changes ranging from 2 − 6% or even up to 10% under physiological loading conditions [63, 39]. Neglecting the compressibility can lead to inaccurate stress predictions, particularly in complex biomechanical simulations where hydrostatic stress states are nonuniform. Consequently, accurate constitutive modeling requires a framework that explicitly accounts for this volumetric variation through the deformation. To incorporate compressibility, Nolan et al. [40] proposed the compressible Holzapfel–Gasser–Ogden (HGO-C) model, which often splits the strain energy into volumetric and isochoric parts, where the anisotropic fiber contribution initially defined for incompressible materials is expressed using isochoric invariants [40]. However, authors have highlighted fundamental limitations with this standard volumetric-isochoric split when applied to anisotropic materials [45, 38]. Specifically, formulating the anisotropic energy component solely in terms of isochoric invariants renders it insensitive to volumetric deformation, potentially leading to physically unrealistic predictions, such as a purely isotropic response under hydrostatic loading. Since anisotropic fibers are expected to contribute to the volumetric response [40, 57], the standard HGO-C formulation can fail to model compressible anisotropic behavior correctly.

To address this deficiency, alternative formulations, such as the modified anisotropic (MA) model proposed by Nolan et al. [40], advocate that the anisotropic strain energy should depend on the invariants of the **full** right Cauchy-Green tensor rather than only its isochoric counterparts to ensure proper coupling between volumetric and anisotropic responses. For this reason, we adopt this updated model, consistent with the MA model principles detailed in Section 2.4 (12). The anisotropic term is formulated using full deformation invariants to capture potential compressible anisotropic effects accurately.

While the HGO framework effectively captures the characteristic stiffening behavior associated with collagen fiber recruitment, applying it involves difficulties beyond simply modeling compressibility. First, many methodologies struggle to robustly account for the known regional variations in the mechanical properties of the aortic wall [32, 44]. Existing approaches often fail to capture the gradual transitions in fiber orientation and arrangement from the ascending to the abdominal aorta, which are essential for understanding the aorta’s integrated mechanical behavior. Second, material parameter identification methods for HGO models often encounter challenges due to the nonconvexity of the optimization landscape, which can lead to convergence to local minima [3]. Additionally, sensitivity to outliers common in experimental biological data can hinder robust parameter estimation. Divergences between model predictions (e.g., for Poisson’s ratios) and experimental data have also been noted, potentially linked to the underlying assumptions about compressibility and anisotropy [53].

### 1.3 Objectives and Current Study

To address the challenges of robustly characterizing regional aortic mechanics, this study aimed to develop and validate a methodology for identifying regional material parameters in the rat aorta. This is achieved using a physically consistent compressible HGO (HGO-C) formulation [40], which explicitly accounts for the gradual transition in properties along the vessel wall. We propose a dual-estimation strategy. First, global optimization identifies the baseline parameters that fit all uniaxial test data from the ring specimens of a given animal from three anatomical regions: the proximal ascending/aortic-arch segment (AOA), the descending thoracic aorta (DTAo), and the descending abdominal aorta (DAAo). Subsequently, the parameters of each anatomical segment were refined by local optimization to accommodate regional variations while maintaining a realistic distribution. This dual estimation approach is designed to capture the smooth transition in elastic properties from the ascending aorta to the descending abdominal aorta more effectively than the conventional single-stage fitting procedures.

Parameter identification from experimental data constitutes an inverse problem, a class of problems that is often ill-posed, potentially leading to non-unique solutions or high sensitivity to measurement noise [13, 58]. To enhance the robustness against experimental outliers and regularize the inverse problem, our optimization framework minimizes a carefully constructed cost function. This function quantifies the misfit between the model predictions and experimental data using the Cauchy loss, which is known for its resilience to outliers, and incorporates a Tikhonov-type regularization term to promote solution stability and uniqueness. The cost function, ℱ, to be minimized is defined as

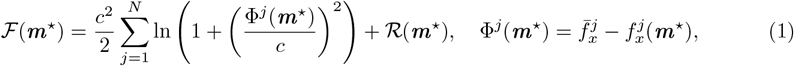

where ℱ denotes the regularized cost function, and the first term quantifies the data misfit between the observed forces 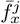 from uniaxial extension tests and the predictions 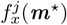 of our variational HGO-C model, using the Cauchy loss for robustness. The residual Φ^*j*^ represents the difference at data point *j*, and *c* is a scaling parameter that modulates the outlier influence. The second term is the regularization penalty, ℛ (***m***^⋆^), weighted by the regularization parameter *ω* ≥0. This term introduces a penalty based on the properties of the parameter set ***m***^⋆^ itself (e.g., its norm or deviation from a baseline estimate), thereby helping to stabilize the solution, select a unique solution from potentially many solutions, and prevent overfitting to noisy experimental data [8, 37].

The overall process involves solving the forward problem (predicting force 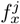 given parameters ***m***^⋆^ using a variational formulation) within an inverse problem framework (finding optimal ***m***^⋆^ to minimize ℱ). The uniaxial extension tests provide records of the reaction forces 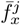 along the x-axis [***e***_1_], performed under quasi-static conditions, where inertial and viscous effects are considered negligible.

To biologically validate the estimated mechanical parameters, uniaxial tensile tests were performed on fresh rings of rat aortas extracted from the arch, thoracic, and abdominal regions. Furthermore, the protein contents of collagen types I and III and elastin were quantified via Western Blot in samples obtained from these regions in a separate cohort of the same animal model to correlate the microstructure with mechanics. Collagen types I and III are the predominant types of collagen present in the aortic wall, accounting for approximately 80 − 90% of the total collagen [4]. Type I collagen primarily provides tensile strength, whereas type III collagen contributes more to the elasticity of the vessel wall.

## 2 Methods

This study employed a single-layer model of the aortic ring, considering samples obtained from the proximal ascending/aortic-arch segment (AOA), descending thoracic aorta (DTAo), and descending abdominal aorta (DAAo) to implement a dual estimation strategy for identifying material parameters. The model conceptualizes the ring as a fiber-reinforced composite material in which stiff collagen fibers are embedded within a homogeneous, isotropic (soft) ground matrix. The inclusion of fiber orientation and dispersion imparts an orthotropic configuration to the material model.

### 2.1 Experimental procedure

All animal experiments were approved by the Institutional Animal Care and Use Committee of the University of São Paulo Medical School (080/16, SDC4398/16/064).

Male Wistar rats (N=5, 8-10 weeks old, 300-350 g) served as subjects for these experiments. Anesthesia was induced with 5.0% isoflurane and maintained with 3.0% isoflurane in *O*2-enriched medical air, ensuring animal comfort throughout the procedure. The animals were placed in the dorsal decubitus position and underwent ultrasonographic examination to assess aortic diameter *ϕ*^*r*^ and wall thickness (*τ* ^*r*^). Images were acquired from three distinct regions: the proximal ascending/aortic-arch segment (AOA), descending thoracic aorta (DTAo), and descending abdominal aorta (DAAo), using a VEVO 2100 system (Visual Sonics, Toronto, ON, Canada) equipped with a 21 MHz transducer. Figure 1 shows representative ultrasonographic images illustrating the determination of the aortic diameter and wall thickness (calculated as the difference between the outer and inner diameters). The mean of three measurements in sections A, B, and C of the same sampled region was used as a representative value for that region and was employed in the analysis. This noninvasive measurement technique ensures that aortic dimensions are assessed under physiological conditions.

**Figure 1:**
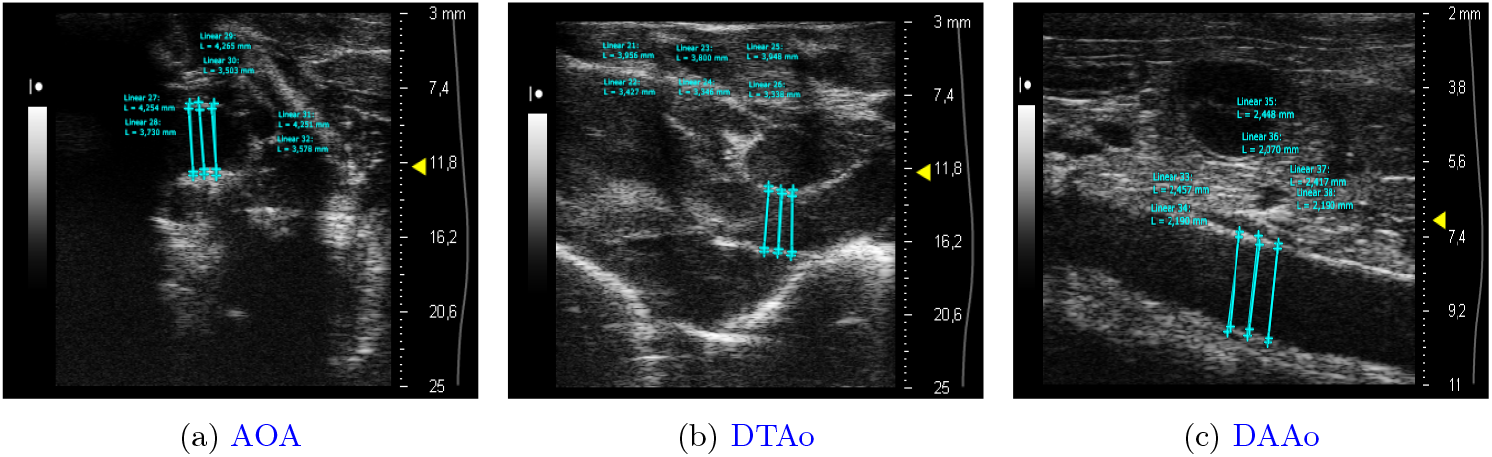
Representative ultrasonographic images showing the determination of aortic wall thickness as the difference between outer and inner diameters in the region of the ascending aortic arch (a), thoracic aorta (b) and abdominal aorta (c).

Following imaging, the carotid artery was cannulated to measure blood pressure (BP) and confirm the normotensive state of the animals. A polyethylene catheter (PE-10) inserted into the artery was connected to a larger bore catheter (PE-50) linked to a pressure transducer (MLT0699; ADInstruments, Bella Vista, NSW, Australia). Arterial blood pressure was recorded continuously for 10 min at a sampling rate of 2 kHz using an analog-digital interface (Power Lab 16/35, ADInstruments, Bella Vista, NSW, Australia). Subsequently, the animals were euthanized with a lethal dose of isoflurane. With the animals in dorsal recumbency, the aorta was exposed and carefully dissected from the heart, and the adjacent connective tissues were removed. The rings were then precisely cut under a microscope from three regions: the proximal ascending/aortic-arch segment (AOA), the descending thoracic aorta (DTAo), and the descending abdominal aorta (DAAo). Specifically, three samples were collected from the proximal segment: one anterior to the brachiocephalic artery trunk and two at the most anterior portion of the aortic arch curvature.

The harvested rings were immediately cooled to 4°*C* in 0.9% NaCl physiological saline solution to maintain tissue integrity and prevent degradation prior to mechanical testing. A second group of animals of similar age and weight (N=9) underwent a comparable procedure to collect aortic samples designated for protein quantification using the Western Blot technique. A detailed description of this biochemical method is provided in Appendix A.

### 2.2 Test protocol

Uniaxial extension tests were conducted using a standard tensile testing machine (Instron©3365, UK) equipped with a temperature-controlled bath (BioPuls Bath, Norwood, MA, USA) to ensure specimen hydration. These tests yielded a series of reaction force records, 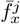, also termed *observations*, along the x-axis [***e***_1_], under quasi-static loading conditions (10.00 mm/min displacement rate), permitting the neglect of inertial and viscous effects. For the classic ring model, ring samples (segments A to C, designated) were extracted from the proximal ascending/aortic-arch segment (AOA), specifically from the region anterior to the brachiocephalic artery branch and the most anterior portion of the aortic arch bend. Additionally, three rings were sectioned from the descending thoracic aorta (DTAo) and three from the descending abdominal aorta (DAAo) between the anterior and posterior mesenteric arteries. This structured regional sampling strategy is crucial for capturing the acknowledged variations in aortic wall properties along its length.

The aortic ring segments were secured in the testing machine using a custom-designed holder featuring U-shaped arms that applied force circumferentially. Each side of the ring was carefully positioned on these arms at ambient temperature before the assembly was mounted on a tensile testing machine. Figures 2a and 2b show the experimental setup. The position of the bath reservoir was adjusted to ensure that each ring was fully immersed in saline solution and maintained at a constant temperature of 37°C (wet condition). Table 1 provides the measured mean thickness (*τ* ^*r*^) and mean diameter (*ϕ*^*r*^) of the aortic rings from the studied anatomical regions. The diameter decreases from the proximal to distal aorta (mean *ϕ*^*r*^: AOA *>* DTAo *>* DAAo), consistent with the known tapering of the aortic lumen along its length. Maintaining the tissue under wet conditions at physiological temperatures is essential for preserving its mechanical integrity and ensuring accurate measurements, which is consistent with the protocols used by Huh et al. [30] and Giudici et al. [20].

**Figure 2:**
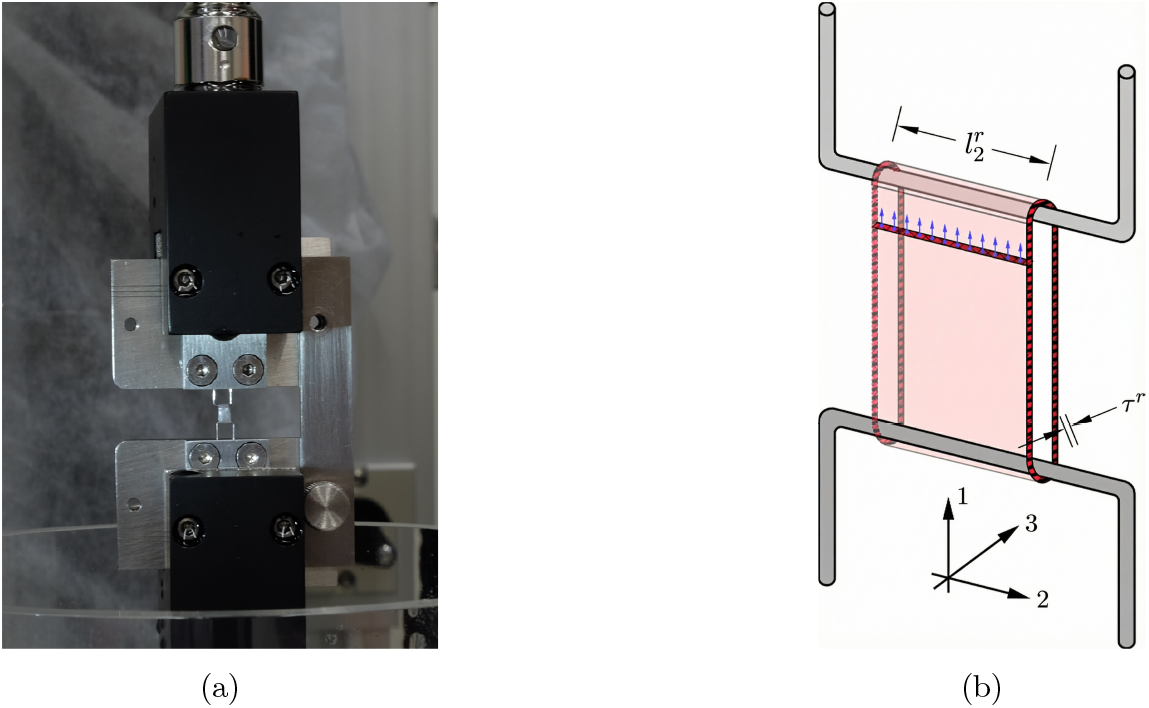
(a) Experimental setup showing the position of the aortic ring in the uniaxial extension tests. (b) Detailed hand-drawn schematic of the experimental setup.

**Table 1:** List of animals studied, with the aorta divided into the proximal ascending/aortic-arch segment (AOA), descending thoracic aorta (DTAo), and descending abdominal aorta (DAAo), the segments (*A, B, C*), and their dimensions (mm): thickness (*τ* ^*r*^), diameter (*ϕ*^*r*^), and width of the ring 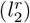.

| Rat<br>N. | Section | $\tau^r$<br>[mm] | $\phi^r$<br>[mm] | Segments width $l_2^r$ [mm] | | |
| --- | --- | --- | --- | --- | --- | --- |
|  |  |  |  | A | B | C |
| 1 | AOA | 0.249 | 3.406 | 1.500 | 1.200 | 1.200 |
|  | DTAo | 0.291 | 3.202 | 1.200 | 1.600 | - |
|  | DAAo | 0.260 | 2.973 | 1.700 | 1.000 | 1.100 |
| 2 | AOA | 0.192 | 3.684 | 1.200 | 1.000 | 1.000 |
|  | DTAo | 0.223 | 3.870 | 1.500 | 1.800 | 1.700 |
|  | DAAo | 0.213 | 2.885 | 1.200 | 1.200 | 1.500 |
| 3 | AOA | 0.349 | 3.243 | 1.200 | 1.000 | 1.200 |
|  | DTAo | 0.350 | 3.195 | 2.000 | 1.800 | 1.600 |
|  | DAAo | 0.238 | 2.960 | 1.500 | 1.500 | 1.800 |
| 4 | AOA | 0.312 | 3.430 | 1.800 | 1.500 | 1.500 |
|  | DTAo | 0.361 | 3.132 | 2.000 | 2.000 | 1.800 |
|  | DAAo | 0.180 | 2.429 | 1.800 | 1.800 | 1.800 |
| 5 | AOA | 0.251 | 2.920 | 1.200 | 1.200 | 1.500 |
|  | DTAo | 0.268 | 2.469 | 1.800 | 1.800 | 1.800 |
|  | DAAo | 0.213 | 2.209 | 2.000 | 1.800 | 2.000 |

The initial separation of the U-shaped arms along the x-axis (pulling direction), denoted 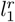, is automatically recorded by the testing machine. The axial width of each ring sample, 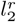(see Table 1), was measured before testing. The testing machine measured the total reaction force 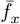 and the displacement *u*_1_ between the U-shaped arms along the x-axis (***e***_1_), which corresponded to the applied ring strain. Each sample underwent preconditioning through three consecutive loading cycles (0.05 mm at 10.00 mm/min) to mitigate the tissue’s hysteresis effect and achieve repeatable force-displacement curves [30]. The force and displacement data acquired during these uniaxial tests were subsequently used to generate the stretch *λ*_*x*_ curves along the ***e***_1_ direction, which served as the prescribed boundary conditions in the forward computational model.

The stretch and engineering stress of the aortic specimen in the direction of the uniaxial extension test are characterized as follows

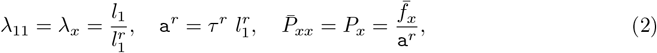

where *λ*_*x*_ is the stretch ratio along the x-axis, 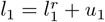 is the current length between the arms in the pulling direction, and 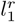 is the initial reference length (i.e., initial arm separation). The term a^*r*^ denotes the initial cross-sectional area of *one* leg of the ring in the reference configuration, calculated from the measured initial thickness *τ* ^*r*^ and initial width 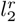. Consequently, 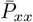 represents the engineering stress in one leg of the ring, computed from the total measured force 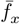, assuming that the load is equally distributed between the two supporting arms. Figure 2 illustrates the schematic setup and the experimental arrangement.

### 2.3 Kinematics

Standard continuum mechanics notation is adopted throughout this work. Scalars are denoted by lightface italic letters (e.g., *J, p*), vectors by boldface lowercase letters (e.g., ***u, n***), and second-order tensors by boldface uppercase letters (e.g., ***F, C***, ***P***)

The ring geometry was modeled as a single arterial wall layer (Fig. 2), represented by a continuum body whose reference placement is denoted by Ω^*r*^ ⊂ ℝ^dim^. This represents a common simplification, in which distinct native aortic layers (intima, media, and adventitia) are homogenized into a single entity with effective material properties. Although this approach enhances model tractability, it inherently averages the specific contributions of each layer. The domain of interest is defined by the material points ***x***^*r*^ in the reference configuration, and their corresponding positions in the current configuration ***x***, linked through a suitably smooth nonlinear deformation map ***φ***: Ω^*r*^ × [0, *T*] → Ω at time *t* ∈ [0, *T*], where *T* ∈ ℝ_+_.

The boundary of the reference configuration Ω^*r*^ is denoted by ∂Ω^*r*^, with an outward unit normal vector ***n***^*r*^. The body is subjected to body forces ***b***^*r*^ per unit volume within Ω^*r*^ and traction forces ***t***^*r*^ = ***Pn***^*r*^ per unit area on a Neumann boundary subset Γ_*N*_ ⊂ ∂Ω^*r*^, where ***P*** is the first Piola–Kirchhoff (PK1) stress tensor. Dirichlet boundary conditions, 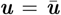, are prescribed on a subset Γ_*D*_ ⊂ ∂Ω^*r*^, specifying the displacement field ***u*** for that portion of the boundary. The motion is characterized by the displacement field ***u***(***x***^*r*^, *t*) = ***φ***(***x***^*r*^, *t*) − ***x***^*r*^.

The map *φ* must belong to the space of kinematically admissible configurations, defined over the Cartesian product of the reference domain and time, thus ℋ := {*φ*: Ω^*r*^ × [0, *T*] → R^*dim*^ | *J*(***x***^*r*^, *t*) := det(∇*φ*(***x***^*r*^, *t*)) *>* 0 in Ω^*r*^ × [0, *T*] and 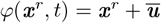 on Γ_*D*_ × [0, *T*}.

### 2.4 Constitutive model

To appropriately handle the quasi-incompressibility characteristic of arterial tissue, the multiplicative decomposition of the deformation gradient ***F*** was employed. This decomposition separates ***F*** into a spherical (dilatational) part, ***F***_vol_ = *J*^1*/*3^***I***, and a unimodular (isochoric or volumepreserving) part, 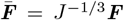, where ***I*** is the second-order identity tensor. This split is particularly advantageous for modeling quasi-incompressible materials where *J* ≈ 1, although the decomposition is general. Based on this decomposition, the right Cauchy-Green tensor ***C*** and its isochoric counterpart 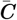 are defined as

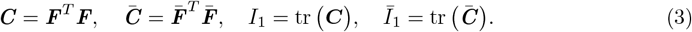

The material is assumed to be hyperelastic, which means its stress response derives from a scalar strain-energy density field *ψ*(***x***^*r*^, *t*). To ensure objectivity, the constitutive counterpart is written as 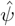 and formulated as a function of ***F*** (or equivalently of ***C***), such that the second Piola–Kirchhoff (PK2) stress is 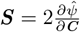 and ***P*** = ***FS***. Accordingly, the stress response emerges directly from the scalar strain-energy functional.

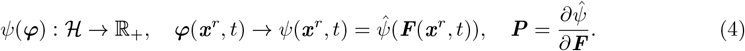

This constitutive potential uses an invariant-based framework for anisotropic materials under finite deformations. The arterial wall is modeled with two families of collagen fibers arranged in a symmetrical spiral, with their preferred directions within the aortic wall’s tangential plane. Collagen stiffening under load is described by an exponential function aligned with these directions, which is a core feature of the HGO model. This approach yields the nonlinear mechanical response of arterial tissue, where the exponential function captures the increase in arterial stiffness as it stretches along the preferred fiber orientation [18, 24, 40, 21].

To outline the phenomenological model, the strain-energy function 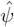, based on the HGO framework, is defined in a decoupled form. This form accounts for the isochoric response of the matrix material (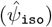), the volumetric response (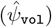), and the anisotropic contribution from the collagen fibers (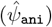)

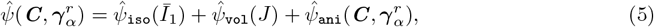

where 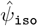 represents the isochoric isotropic contribution of the non-collagenous ground matrix (dependent on the first isochoric invariant *Ī*_1_), 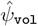 signifies the volumetric contribution (dependent on the Jacobian *J*), and 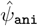 captures the anisotropic contribution arising from two families of collagen fibers (dependent on the full deformation tensor ***C*** and the fiber direction vectors 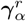). The fiber families are assumed to be symmetrically oriented with respect to the Cartesian axes 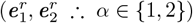. Their preferred directions in the reference configuration are defined by unit vectors 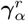

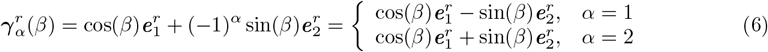

Here, *β* ∈ [−*π/*2, *π/*2] is the preferred Eulerian angle of each collagen fiber family relative to the principal material axis, with 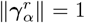. Associated with each 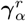, the pseudo-invariants of ***C*** for each fiber family are defined as

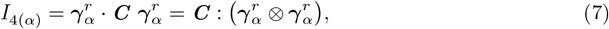

where 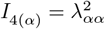 represents the square of the stretch (*λ*_*αα*_, *α* ∈ {1, 2}) along the direction of the *α*-th collagen fiber family. This invariant is generally sufficient to capture the typical anisotropic features observed in arterial wall responses [23] and is commonly used in models of fiber-reinforced hyperelastic materials [12].

To account for the inherent non-uniform orientation of the collagen fibers, a dispersion factor was incorporated into the model. This approach, introduced by Gasser et al. [18], acknowledges that while collagen fibers have a predominant orientation along 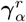, they are not perfectly aligned but rather exhibit a preferred yet dispersed orientation. Incorporating a dispersion factor as a scalar structural parameter, *κ*, allows for a more realistic representation of the elastic behavior of the arterial wall. We utilize a generalized structural tensor 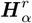, which represents a transversely isotropic distribution of fibers symmetrically dispersed around the mean direction 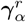. The structural tensor 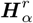 is defined as

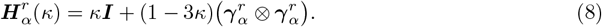

The parameter *κ* ∈ [0, 1*/*3] quantifies the degree of anisotropy arising from this fiber dispersion and is typically characterized by experimental data on fiber orientation distributions [18]. An isotropic distribution of fiber orientations (maximum dispersion) corresponds to *κ* = 1*/*3, whereas *κ* = 0 indicates the perfect alignment of fibers (no dispersion). By combining the right Cauchy-Green tensor ***C*** with the structural tensor 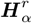, modified pseudo-invariants 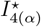 are obtained

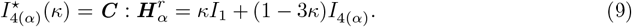

These modified pseudo-invariants 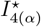 significantly influence the strain-energy density function under fiber tension. Their contribution is considered negligible under compression, as fibers typically cannot support compressive loads. In the uniaxial tensile test setup, the aortic ring experiences only tensile loads. Therefore, compressive fiber behavior is not explicitly modeled. Details on filtering mechanisms for tensile-only deformations are available in Federico and Gasser [14]. Other approaches employ continuous probability density distributions (e.g., bivariate von Mises or Bingham distributions) integrated over the unit sphere (Angular Integration (AI) methods) to model angular fiber dispersion [47, 27]. Instead, we adopted the generalized structural tensor (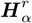) and dispersion parameter (*κ*) proposed by Gasser et al. [18]. Comparative studies and reviews indicate that, for rotationally symmetric dispersions, the macroscopic mechanical responses predicted by the Generalized Structural Tensor (GST) framework closely approximate those obtained from full angular integration, particularly in low-dispersion regimes. However, angular-integration models can explicitly represent a broader spectrum of fiber distributions and may be preferable when it is necessary to resolve wider dispersion states. In this study, the fitted solutions were concentrated near low dispersion; therefore, the GST model was sufficient for inverse analysis, maintaining computational efficiency and tractability during parameter estimation. [25, 28, 15].

The isochoric isotropic response (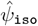) of the ground matrix is modeled using the neo-Hookean form [12], which depends on the first isochoric invariant *Ī*_1_

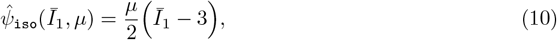

where *µ >* 0 is the shear modulus of the ground matrix material.

Given the experimental evidence suggesting that arterial walls exhibit slight compressibility [63, 39, 9], a compressible model was employed. Consistent with the corrected HGO (HGO-C) approach by Nolan et al. [40], the volumetric strain-energy function (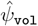) is defined as dependent on the Jacobian *J* [50] as follows

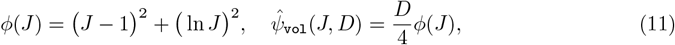

where *D >* 0 denotes the bulk modulus (or penalty parameter) and represents the material’s resistance to volumetric changes. Crucially, adhering to the Modified Anisotropic (MA) correction principles [40], the anisotropic strain-energy term 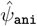 depends on the **full** right Cauchy-Green deformation tensor ***C*** (via the invariants *I*_4(*α*)_), rather than solely on its isochoric part 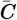. Consequently, the anisotropic part of the strain-energy density function, 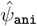, is expressed as

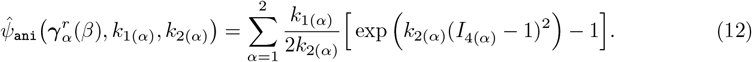

In this expression, *k*_1_ *>* 0 is a stress-like parameter linked to the stiffness of the collagen fibers, and *k*_2_ *>* 0 is a dimensionless parameter that characterizes the nonlinearity of the collagen fiber response. To incorporate the effects of dispersion, the dispersion factor *κ* (by using the modified invariant 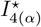 from (9)) is introduced into the anisotropic energy expression (12). With this modification, the resulting dispersed anisotropic strain-energy density 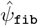 is given by

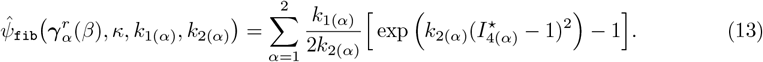

For simplicity, we assumed that the three layers of the arterial wall were effectively lumped into one unit, consisting primarily of two families of symmetrically arranged collagen fibers. The material parameters for both fiber families are assumed to be identical and are summarized as ***m***_fib_ = {*k*_1_, *k*_2_, *β, κ*}. Considering the isochoric isotropic (*µ*), volumetric (*D*), and fiber contributions, the complete set of material parameters for the model is defined as

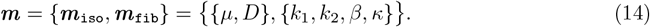

### 2.5 Three-field variational formulation

To numerically handle the near-incompressibility constraint arising from the volumetric term 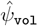, a mixed variational approach based on the extended potential energy of Hu-Washizu functionals [29, 51] is employed. This method is formulated as a three-field problem comprising the displacement field ***u*** ∈ 𝒱, an independent dilatation field Θ ∈ ℒ, and an independent hydrostatic pressure field *p* ∈ ℒ. Treating these fields as independent kinematic variables enhances the flexibility and stability of the numerical solution, particularly for quasi-incompressible materials undergoing finite deformations [52, 33].

The function space ℒ for the fields Θ and *p* is defined as ℒ:= {*q* ℒ^*h*^(Ω^*r*^)}, representing square-integrable functions on Ω^*r*^ (typically *L*^2^(Ω^*r*^) after discretization, where *h* relates to element size). The three-field functional Π: 𝒱 × ℒ_Θ_ × ℒ_*p*_ → ℝ_+_ is given by the expression

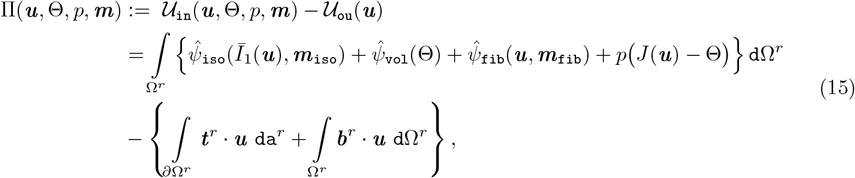

where Θ represents a volume change measure independent of the motion, while *p* acts as a Lagrange multiplier field enforcing the local volumetric constraint *J*(***u***) = Θ. The flexibility afforded by using independent variables for ***u***, Θ, *p* helps to prevent volumetric locking phenomena in numerical implementations [48].

### 2.6 Balance equations

The principle of virtual work follows from enforcing the stationarity of the Hu-Washizu functional Π in equation (15). This principle must also include additional terms for the quasi-incompressibility constraints. Taking independent variations of Π with respect to the displacement ***u***, dilatation Θ, and hydrostatic pressure *p* yields the weak forms of the governing equations [7, 62].

The space of kinematically admissible variations, or the tangent space to ℋ, is denoted by 𝒱. Let *δ****u*** ∈ 𝒱_0_, and *δ*Θ, *δp* ∈ ℒ be admissible test functions, where V_0_ := {*δ****u*** ∈ [*H*^1^(Ω^*r*^)]^3^ |*δ****u*** = ***0*** on Γ_*D*_}. The variations of Π with respect to ***u***, Θ, and *p* yield the following system of equations by find ***u***, Θ, *p* ∈ {𝒱 × ℒ_Θ_ × ℒ_*p*_} such that for all (*δ****u***, *δ*Θ, *δp*) ∈ {𝒱_0_ × ℒ_Θ_ × ℒ_*p*_}

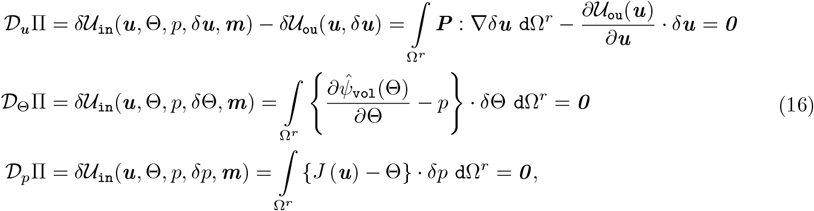

To solve the equilibrium condition given by the first expression in equation (16), a constitutive relation for the PK1 stress tensor ***P*** is needed. This tensor is derived from the total strain-energy function 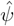(5), and the relation follows the standard hyperelastic formulation

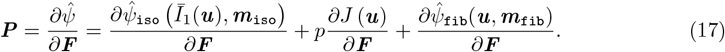

The first equation in (16) represents the equilibrium condition, or the virtual work statement, while the second equation establishes the constitutive relation for the pressure field by linking the pressure to the derivative of the volumetric strain-energy function 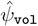 with respect to the independent dilatation Θ, and the third equation enforces the volumetric constraint *J*(***u***) = Θ in a weak sense. Notably, alternative sign conventions for the hydrostatic pressure *p* exist in the broader mechanics literature. By defining the volumetric stress contribution as 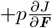, we follow the standard solid mechanics convention for mixed variational formulations [7, 62], in which a positive *p* corresponds to a tensile hydrostatic stress.

By solving this coupled system of equations (16), we obtain the displacement field ***u***, the dilatation field Θ, and the hydrostatic pressure field *p*. Together, these fields describe the mechanical response of the aortic wall under applied loads and boundary conditions.

### 2.7 Simple tension

In a uniaxial tensile test, the aortic ring is subjected to strictly macroscopic one-dimensional loading. Assuming the tissue is tested along its circumferential direction, with the loading axis aligned to the spatial *x*-direction. The deformation gradient ***F*** in this locally aligned coordinate system is a purely diagonal tensor, mapping the reference material frame (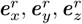) to the spatial current frame (***e***_*x*_, ***e***_*y*_, ***e***_*z*_).

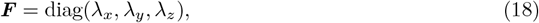

Here, *λ*_*x*_ is the primary stretch along the loading direction, while *λ*_*y*_ and *λ*_*z*_ are the principal transverse stretches corresponding to radial thickness and axial width. The right Cauchy-Green tensor is 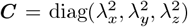, and the volume ratio is *J* = *λ*_*x*_*λ*_*y*_*λ*_*z*_. Under this diagonal deformation, the modified first principal invariant *Ī*_1_ = *J*^−2*/*3^ tr(***C***) expands explicitly to

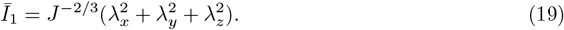

Likewise, modified pseudo-invariants like 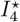, which govern the anisotropic response of collagen fibers, can be expressed as scalar functions of the principal stretches and the fibers orientation angles. Since the deformation is purely normal and the coordinate frames are aligned, the PK1 stress tensor ***P*** is strictly diagonal. Although ***P*** is a two-point tensor relating the reference and current configurations, the aligned bases allow us to use principal nominal stresses without loss of rigor. The diagonal tensor 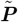 holds the principal normal stresses of ***P***, represented as

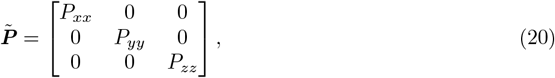

where each principal stress components *P*_*xx*_, *P*_*yy*_, *P*_*zz*_ are obtained by taking the derivative of the total strain-energy function 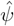 with respect to the corresponding stretch *λ*_*i*_. Utilizing the mixed formulation introduced in Section 2.6, the principal stresses have this explicit analytical form

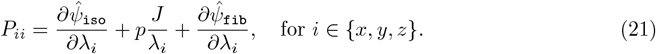

During uniaxial extension, the surfaces of the ring normal to the y and z directions are traction-free. We simplify the analysis by adopting the plane stress hypothesis, which is common for thin biological membranes and shells. This assumption posits that stresses acting perpendicular to the ring plane (*P*_*zz*_ and shear stresses *P*_*xz*_, *P*_*yz*_) are negligible compared to in-plane components (*P*_*xx*_, *P*_*yy*_, *P*_*xy*_). Though approximate, this assumption reduces dimensionality and simplifies the problem. It is justified by

- **The geometry**: The aortic ring specimens are thin relative to their diameter and width.
- **The loading**: The primary load is applied circumferentially, predominantly within the plane of the ring.
- **Expected behavior**: For thin structures subjected to in-plane loading, through-thickness stresses are generally small.

Under this assumption, the transverse principal stress components must vanish (*P*_*yy*_ = 0 and *P*_*zz*_ = 0). Consequently, the only non-zero component of the stress tensor balancing the externally applied load is the primary uniaxial stress, *P*_*xx*_. This theoretical stress is directly related to the experimentally measured uniaxial force introduced in equation (2). By enforcing plane stress conditions alongside the volumetric soft constraints (quasi-incompressibility) from the mixed variational framework, we obtain a coupled nonlinear system of equations. Solving this system yields the unknown transverse stretches (*λ*_*y*_, *λ*_*z*_) and hydrostatic pressure *p*, all as functions of the primary driving stretch (*λ*_*x*_). The resulting predicted uniaxial force can then be directly evaluated and compared against the experimental data to identify the optimal material parameters.

## 3 Material Parameter Identification

The determination of material parameters (***m***) for the constitutive model from experimental data is an inverse problem. This section outlines our methodological approach, which integrates a robust cost function, tailored regularization strategy, efficient gradient computation via the adjoint method, and advanced optimization algorithms. These components collectively address the challenges inherent in fitting complex nonlinear models to biological data, which are often characterized by noise and outliers.

A central feature of our methodology is the dual estimation strategy, which is detailed in Section 3.5. This strategy is based on the hypothesis that the aortic microstructure exhibits a physiologically smooth transition along its length. It aims to mitigate issues such as overfitting and reduce the variability often observed in the estimated parameters when using traditional fitting techniques.

### 3.1 Forward Problem Solution

The forward problem involves simulating a uniaxial ring extension test using the previously developed variational formulation. Given a candidate set of material parameters ***m***, the stationarity conditions derived from the principle of virtual work, as stated in equation (16), yield a system of nonlinear algebraic equations. This system is solved iteratively, by employing a quasi-Newton method for each prescribed stretch level *j* from the experimental data. The solution of this system provides the state variables ***v***^*j*^ = {***u***^*j*^, Θ^*j*^, *p*^*j*^} (which include displacements ***u***^*j*^, the independent dilatation field Θ^*j*^, and the hydrostatic pressure field *p*^*j*^). These computed state variables subsequently enable the calculation of the predicted reaction forces 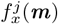 based on the input set of material parameters ***m***. This predicted force is expressed as

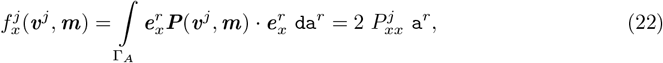

where ***P***^*j*^ and its relevant component 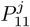 are the PK1 stress tensor (17) and its component in the loading direction, respectively, evaluated at the *j*-th stretch level. Γ_*A*_ represents the cross-sectional area of one leg of the ring sample, and 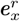 is a unit vector in the loading direction.

### 3.2 Inverse Problem Formulation and Regularization

The inverse problem aims to identify the optimal set of material parameters ***m***^⋆^ that minimizes the cost function 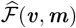. This function quantifies the misfit between the model predictions, *f*_x_ (***v, m***), and the experimentally observed data, 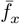, subject to feasible bounds ***m***_min_ ≤ ***m*** ≤ ***m***_max_.

Parameter identification from experimental data, especially biological data, is susceptible to several well-known issues

- **Ill-posedness**: Inverse problems are frequently ill-posed, meaning that minor errors or noise in the input data (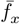) can lead to significant variations in the estimated parameters (***m***). The solutions may not be unique or may not depend continuously on the provided data [13, 58].
- **Overfitting**: Complex models endowed with numerous parameters can fit the calibration data (including inherent noise) exceptionally well but may consequently fail to generalize the behavior under different conditions. This is particularly important when utilizing potentially noisy experimental data [17, 49].
- **Non-convexity**: The cost function landscape typically exhibits multiple local minima, making it challenging for standard optimization algorithms to locate the global optimum [3].
- **Outliers**: Biological data usually contain outliers arising from experimental artifacts or inherent sample variability. Standard least-squares cost functions are notably sensitive to these outliers.

Our methodology employs several strategies to address these issues. To enhance robustness against outliers and to provide a suitable measure of data misfit within a regularized framework, we define the cost function using the Cauchy loss combined with regularization terms

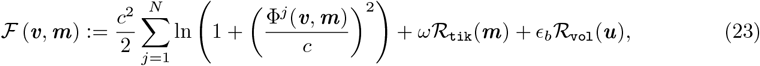

where 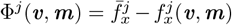 is the residual between the *j*-th experimental observation and the model prediction.

- The **Cauchy loss** term, characterized by the scaling parameter *c*, reduces the influence of large residuals (potential outliers), thereby improving fitting stability. The choice of *c* tunes the robustness; a smaller *c* makes the loss more robust to large errors, whereas a larger *c* approaches the L2 loss behavior. While the underlying constitutive model (HGO-C) builds on the corrections suggested by Nolan et al. [40], the use of Cauchy loss is an independent methodological choice made in this study to improve the stability of the fitting process.
- **Tikhonov regularization** (ℛ_tik_(***m***)), weighted by a regularization parameter *ω* ≥ 0, penalizes deviations of the current parameter set ***m*** from a reference parameter set 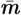

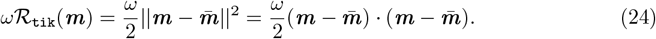

In the first stage of our dual estimation (Section 3.5), 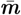 often represents an initial estimate (e.g., zero or parameters close to ***m***_min_). In contrast, the second stage is a set of baseline parameters derived from the first stage. This term mitigates ill-posedness by incorporating prior information, promoting smoother solutions, and aiding the selection of a unique, stable solution from potentially many [13, 8].
- The **Volume regularization** term, ℛ_vol_(***u***), is introduced to address the challenge of identifying the bulk modulus *D* (and thus the volumetric behavior) from uniaxial testing. Soft biological tissues have often been idealized as strictly incompressible; recent experimental evidence [39, 63] demonstrates that arterial walls exhibit measurable compressibility: typically 2–10% volume change under physiological loads. Adopting a quasi-incompressible formulation captures this physiological reality. It also robustly mitigates numerical pathologies, such as volumetric locking, in standard finite element implementations. This approach renders inverse parameter estimation ill-posed when only 1D uniaxial data are used. In this setup, transverse strains are not measured, so an unconstrained optimization algorithm can exploit relaxed volumetric kinematics to minimize the axial force residual artificially. This may result in non-physical tissue swelling or shrinkage (*J* ≫ 1 or *J* ≪ 1). To resolve this non-uniqueness, we propose a regularization term. It is based on the volumetric strain energy function component *ϕ* from (11), summed over the *N* data points:

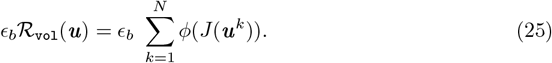

Here, *J* is the Jacobian (volume ratio) computed from the forward solution at the *k*-th data point using the displacements ***u***^*k*^. By enforcing physiological near-incompressibility (*J* ≈ 1) directly within the objective function as a regularization precaution, we appropriately constrain the parameter space. This approach acts as a physics-based soft constraint rather than fixing *D* to an arbitrary constant, thereby mitigating the risk of identifying non-physical parameter sets. It guides the model to properly fit the macroscopic 1D mechanics while maintaining volume changes within physically plausible bounds for a slightly compressible arterial wall. The volumetric regularization weight *ϵ*_*b*_ = 10^−3^ was selected empirically by testing a range of weights across several orders of magnitude and verifying that the resulting Jacobian *J* remained within 2–10% volume change, consistent with aortic tissue measurements [39, 63]. Because uniaxial force-stretch data lack direct information on lateral deformation, the optimizer does not identify a proper gradient information for *D* as a bulk modulus; instead, quasi-incompressibility is assisted via the *ϵ*_*b*_ ℛ_vol_(***u***) term. Thus, *D* values in Table 3 are regularization-influenced descriptors ensuring thermodynamic consistency rather than direct estimates of the physical bulk modulus. Changing *ϵ*_*b*_ by an order of magnitude would shift the fitted *D* and alter the volumetric constraint on *J* during optimization through less or more incompressible behavior. The sensitivity of the identified *D* to *ϵ*_*b*_ is a limitation identified here. Here, *ϵ*_*b*_ denotes the baseline regularization weight used in the first estimation stage (Step 1); the local refinement (Step 2, Section 3.5) uses a distinct weight *ϵ*_*r*_.

The regularization parameters *ω* and *ϵ*_*b*_ serve to balance the adherence to the experimental data against the strength of regularization functions.

### 3.3 Sensitivity Analysis

Gradient-based optimization algorithms, as discussed in Section 3.4, require the computation of the gradient of the cost function ℱ with respect to the material parameters ***m***. This gradient, 𝒟_***m***_ ℱ, comprises contributions from the Cauchy loss and regularization terms. While the gradients of the regularization terms (ℛ_tik_(***m***) and ℛ_vol_(***u***)) can typically be computed directly from their definitions, the gradient of the Cauchy loss term requires the sensitivity of the model output (specifically, the predicted force 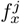) with respect to the parameters ***m***, that is, the derivative 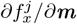.

Directly computing this sensitivity (e.g., via finite differences or analytical differentiation through the entire forward model chain) can be computationally prohibitive for complex nonlinear models involving a large number of parameters. Therefore, we employ the efficient **adjoint method** [58] to compute the required gradient 𝒟_***m***_ ℱ.

The fundamental idea of the adjoint method is to circumvent the explicit computation of the sensitivity of the state variables (such as displacements ***u***, dilatation Θ, and pressure *p*) with respect to each material parameter ***m***_*k*_. Instead, it introduces a set of auxiliary variables known as adjoint variables **Λ**. These are computed by solving a single linear system derived from the governing state equations. Let the state equations (represented by the weak form given in (16)) be denoted as *δ*Π(***v***(***m***), ***m***) = ***0***. The cost function component of interest is 𝒥 (***v***(***m***), ***m***) (here, 𝒥 specifically represents the Cauchy loss part of the total cost function ℱ or an arbitrary function).

The total derivative of 𝒥 with respect to ***m*** is

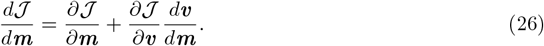

From the state equation, implicit differentiation yields 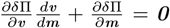, which leads to 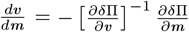. Substituting this back into the expression for the total derivative gives

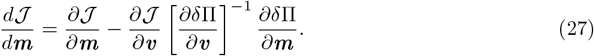

The adjoint method elegantly avoids the computationally expensive inversion of the Jacobian matrix ∂*δ*Π*/*∂***v*** by defining the adjoint variable vector **Λ** as the solution to the linear adjoint equation

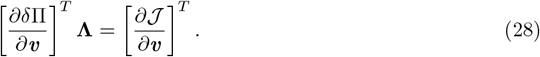

This allows the gradient to be rewritten in a more computationally tractable form

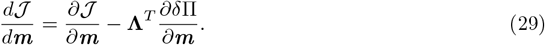

This final form requires only the direct partial derivatives of the cost function 𝒥 and the residuals of the state equations *δ*Π with respect to the parameters ***m***, along with the solution of one linear adjoint system (28) per evaluation of the gradient of the cost function. This is highly efficient, because its cost is independent of the number of parameters in ***m***. The analytical derivations required for the terms ∂𝒥*/*∂***v***, ∂*δ*Π*/*∂***v***, and ∂*δ*Π*/*∂***m*** stem directly from the variational formulation and constitutive laws. Specific expressions for the resulting derivatives of the reaction force *f* ^*j*^ are provided in Appendix B.

### 3.4 Optimization Algorithm

The optimization task entails finding the parameter set ***m*** that minimizes the regularized cost function ℱ, as defined in Equation (23), subject to the bound constraints specified in (30). Owing to the inherent nonlinearity of the constitutive model and the potential for multiple local minima in the cost function landscape [3], a robust optimization strategy is required.

We utilize the Basin-Hopping (BH) algorithm [60] for both the baseline parameter estimation (detailed in Section 3.5, Step 1) and the subsequent local parameter refinement (Section 3.5, Step 2). BH is a global optimization metaheuristic specifically designed to explore complex energy or cost landscapes. It operates by iteratively performing the following sequence of actions

1. Perturbing the current best-known solution randomly to generate a new starting point.
2. Performing a deterministic local minimization search commencing from this perturbed point.
3. Accepting or rejecting the new minimum found based on a probabilistic criterion (e.g., the Metropolis criterion), which allows for occasional uphill moves, enabling the algorithm to escape local basins of attraction.

This strategic combination of stochastic global exploration and deterministic local refinement renders the BH particularly effective at locating global or near-global optima for challenging non-convex optimization problems [35, 42].

For the critical **local minimization** step embedded within each BH iteration, we employ the Interior Point OPTimizer (IPOPT) [6]. IPOPT is a powerful gradient-based algorithm well-suited for large-scale nonlinear programming (NLP) problems, especially those involving bound constraints (such as our ***m***_min_ ≤ ***m*** ≤ ***m***_max_). It incorporates advanced techniques such as interior-point methods and line searches to handle constraints and navigate optimal solutions. Our implementation efficiently leverages the gradients computed using the adjoint method (Section 3.3) to supply first-order derivative information to IPOPT. IPOPT then internally approximates second-order information, frequently utilizing algorithms such as the Limited-memory Broyden-Fletcher-Goldfarb-Shanno (L-BFGS) method [65] for its quasi-Newton updates. This approach enhances the convergence speed without necessitating the explicit computation and storage of the Hessian matrix.

Thus, we address the problem of global minimization of a cost function ℱ, assumed to be of class *C*^2^. The parameters, denoted ***m*** ∈ ℝ_+_, are restricted to positive real values. In this parameter space, the function ℱ may have multiple local minima, ℱ (***m***^*k*^), where each ℱ (***m***^*k*^) corresponds to a local minimizer ***m***^*k*^. The global search algorithm seeks a solution ***m***^∗^ that improves upon all previously encountered local minimizers, i.e., ℱ (***m***^*k*^) ≤ ℱ (***m***^*k*−1^) for all *k*. The optimization problem is formally stated as

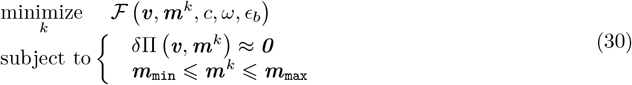

where *δ*Π ≈ ***0*** represents the satisfaction of the stationarity conditions (equilibrium equations) (16), and the inequality constraints define the feasible domain for the material parameters ***m***.

### 3.5 Data Fitting with Double Estimation

We present a dual-estimation strategy to address limitations of traditional material-parameter fitting methods for aortic tissue. The approach assumes that regional properties vary gradually along the aorta and uses regularization to estimate smooth subject-specific trends.

Our methodology was based on the hypothesis that the aortic microstructure exhibits a gradual transition from the proximal ascending/aortic-arch segment to the descending abdominal aorta. These smooth trends should be interpreted as model-guided regularized estimates rather than direct proof of continuous biological transitions.

The dual estimation strategy consists of two sequential steps:

1. **Baseline Parameter Estimation** In the initial material parameters estimation step, we determined a common set of *baseline material parameters* (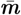) that best represented the shared underlying mechanical characteristics of the aorta within a single animal. These parameters aim to reflect potential similarities before regional adaptations are considered, effectively representing the typical or average behavior across all ring samples from an individual. This methodology involves identifying the parameter set that minimizes the *sum* of the regularized cost functions, as defined in equation (23), across all aortic segments sampled from that particular animal

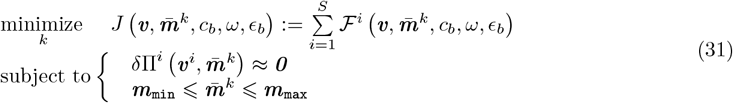

Here, ℱ^*i*^ is the regularized cost function for the *i*-th ring sample, evaluated with a common set of baseline parameters, 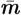. The state equations *δ*Π^*i*^ ≈ ***0*** enforce equilibrium for each sample *I* at the chosen 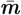. In the global stage, the Tikhonov term 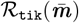 penalizes deviation of 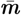 from a predefined initial estimate. This initial estimate could be zero or parameter values near the lower bounds ***m***_min_). The regularization weight *ω* = 10^−4^ determines the strength of this penalty, providing 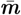 flexibility within the admissible parameter space. The volume regularization term ℛ_vol_(***u***) is applied to each sample to discourage unphysical volume changes, weighted by *ϵ*_*b*_ = 10^−3^ as defined in Equation (25). The Cauchy scaling parameter *c*_*b*_ controls the influence of large residuals in ℱ^*i*^ and was fixed at 40 (after empirical tests over 1 ≤ *c* ≤ 100) to balance robustness and sensitivity to dominant trends. The choice of *c*_*b*_ led to optimized baseline parameters 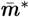, which capture the subject-specific mechanical signature of the aorta and provide the reference solution for later refinement. A global optimization algorithm, BH, used IPOPT as its local solver to efficiently explore the parameter space and overcome suboptimal minima.
2. **Local Parameter Refinement** In the second estimation step, we refined the material parameters ***m***^*i*^ for each ring sample *i* to capture its specific regional mechanical properties. We used the globally optimized baseline parameters 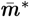 obtained in the first step as the *reference* point for the Tikhonov regularization term (24) in this local stage. The optimization problem for the *i*-th ring sample is then formulated as finding the specific ***m***^*i*^ that minimizes

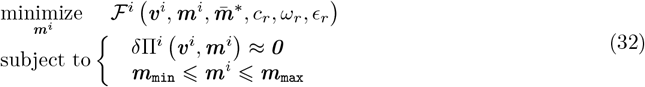

Here, ℱ^*i*^ is again the regularized cost function from (23), minimized over sample-specific parameters ***m***^*i*^. The Tikhonov term penalizes deviation from the baseline 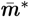, weighted by a regularization parameter *ω*_*r*_ (e.g., *ω*_*r*_ = 10^−3^). The volume regularization term ℛ_vol_(***u***) is also included in Equation (32). This strategy imposes soft constraints by encouraging solutions near the baseline, while not enforcing strict homogeneity. The Cauchy scaling parameter *c*_*r*_ can be adjusted locally (e.g., *c*_*b*_ = 40, *c*_*r*_ = 40); however, in this study, it was kept fixed (*c*_*r*_ = 40) during BH iterations. Increasing *c*_*r*_ relative to *c*_*b*_ allows greater deviation before the residual influence declines, thus broadening the region where the Cauchy loss approximates least squares and balancing sensitivity to sample-specific data with reduced outlier effects. The BH algorithm, using IPOPT locally, robustly solves the nonconvex local refinement problem. Here, *ω*_*r*_ = 10^−3^ stabilized the search around 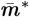. This regularization attracts the search toward the baseline, but does not restrict it: if local data indicates a distinct optimum, the data-misfit term drives the solution elsewhere. This formulation thus balances global consistency (via 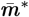) with local flexibility, so segment-specific parameters are regularized estimates of heterogeneity, not unconstrained reconstructions.

## 4 Results

This section presents the principal findings of the study, commencing with the experimental results derived from the uniaxial tension tests and protein quantification assays. Subsequently, the outcomes of the material parameter identification process achieved using the dual-estimation strategy are detailed. An analysis of the resulting stress distributions within the modeled aortic segments follows.

### 4.1 Protein Quantification Across Aortic Segments

The contents of Elastin, Collagen Type I (Col 1), and Collagen Type III (Col 3) were quantified in aortic segments collected from a cohort of control animals (N=9) using the Western Blot technique. Arbitrary Units (AU) are reported following normalization to a consistent loading control (e.g., GAPDH or total protein stain) to ensure comparability across samples and gels. The results are presented in Table 2 along with the P-value from a One-Way ANOVA test. The relative distributions of these proteins across the proximal ascending/aortic-arch segment (AOA), descending thoracic aorta (DTAo), and descending abdominal aorta (DAAo) are shown in Figure 3.

**Table 2:**
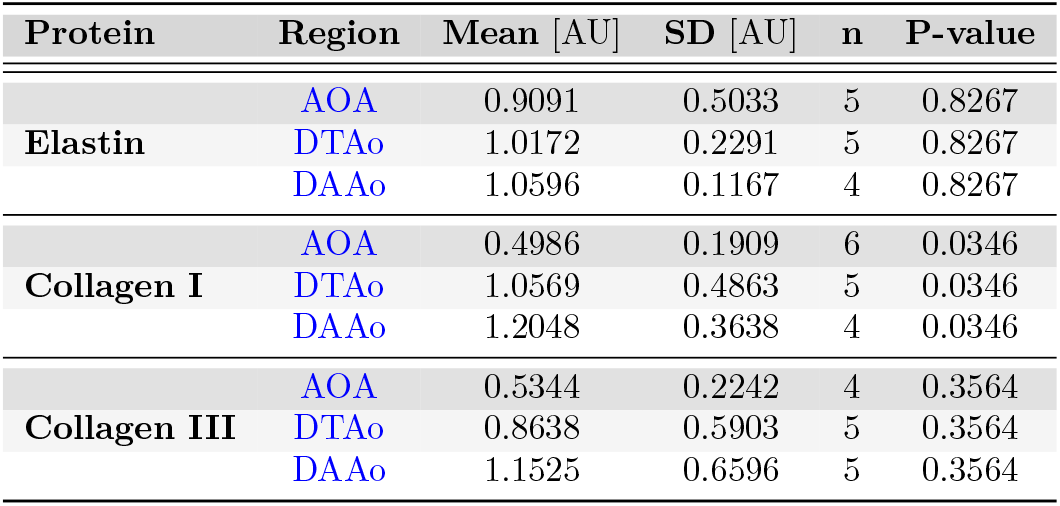
Summary of relative protein quantification (Mean and Standard Deviation, SD) in aortic segments of WT rats, AU = Arbitrary Units, and n indicates the number of samples analyzed for each group.

| Protein | Region | Mean [AU] | SD [AU] | n | P-value |
| --- | --- | --- | --- | --- | --- |
| Elastin | AOA | 0.9091 | 0.5033 | 5 | 0.8267 |
|  | DTAo | 1.0172 | 0.2291 | 5 | 0.8267 |
|  | DAAo | 1.0596 | 0.1167 | 4 | 0.8267 |
| Collagen I | AOA | 0.4986 | 0.1909 | 6 | 0.0346 |
|  | DTAo | 1.0569 | 0.4863 | 5 | 0.0346 |
|  | DAAo | 1.2048 | 0.3638 | 4 | 0.0346 |
| Collagen III | AOA | 0.5344 | 0.2242 | 4 | 0.3564 |
|  | DTAo | 0.8638 | 0.5903 | 5 | 0.3564 |
|  | DAAo | 1.1525 | 0.6596 | 5 | 0.3564 |

**Figure 3:**
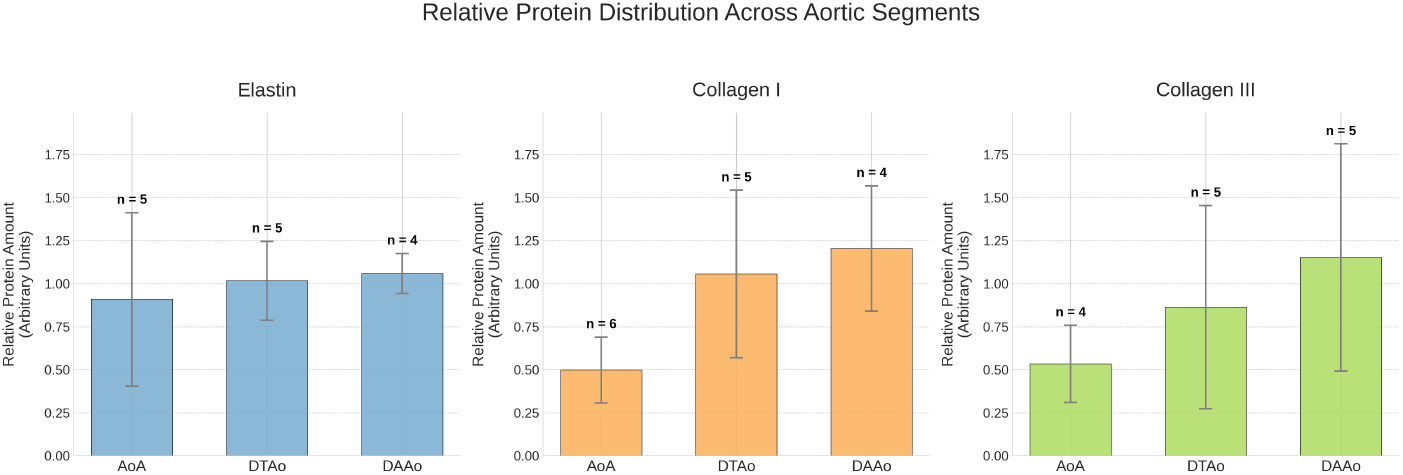
Relative amounts of elastin, collagen I, and collagen III in the proximal ascending/aortic-arch segment (AOA), Descending Thoracic Aorta (DTAo), and Descending Abdominal Aorta (DAAo) segments of WT rats, AU = Arbitrary Units, and n indicates the number of samples analyzed for each region.

The quantification revealed a distinct distribution pattern for these major structural proteins along the aorta, as illustrated in Figure 3 and detailed in Table 2. Despite a similar content of **Elastin** in samples of the three regions studied, there is a trend of its content in the ascending aorta and arch being larger than that of Col type 1 and Col type 3. This relative abundance of elastin throughout the aorta underscores its fundamental role in providing baseline arterial elasticity [59, 56], which contributes to the Windkessel effect [61].

The content of **Collagen Type I** was larger in the DAAo than in the AOA, which is in agreement with its role as a load-bearing element and the increased structural stiffness of the distal aorta [31, 56]. The content of **Collagen Type III**, which contributes to compliance, showed a trend of increasing from the proximal to the distal end.

Despite the limited sample size and inherent biological variability, the observed trends in regional differences are in agreement with the well-described interplay between regional composition and tensile mechanical behavior (Section 4.2).

### 4.2 Force-Stretch Curves

Figure 4 shows the force-stretch curves from uniaxial extension tests of aortic rings of the proximal ascending/aortic-arch segment (AOA), descending thoracic aorta (DTAo), and descending abdominal aorta (DAAo) in 5 rats. The corresponding fitted curves (continuous lines) from the constitutive model are shown along with the force-stretch curves. The curves consistently exhibit the characteristic nonlinear mechanical response of biological tissues, often referred to as *pseudoelasticity* [16, 2]. As established in the arterial mechanics literature (e.g., Holzapfel and Ogden [24]), soft biological tissues are inherently viscoelastic and show history-dependent behavior. This behavior includes hysteresis and internal energy dissipation. After appropriate experimental pre-conditioning using repeated cyclic loading, the stress-strain response becomes steady and highly repeatable. The concept of ‘pseudoelasticity’ acknowledges this preconditioned state. This provides theoretical justification to approximate complex, dissipative tissue behavior using purely hyperelastic strain-energy functions evaluated along a specific, consistent loading path. The initial response reflects the behavior of highly extensible elastin fibers. This is followed by an exponential increase in stiffness with stretch ratio (*λ*_*x*_), due to much stiffer, initially crimped or undulated collagen fibers [59, 56].

**Figure 4:**
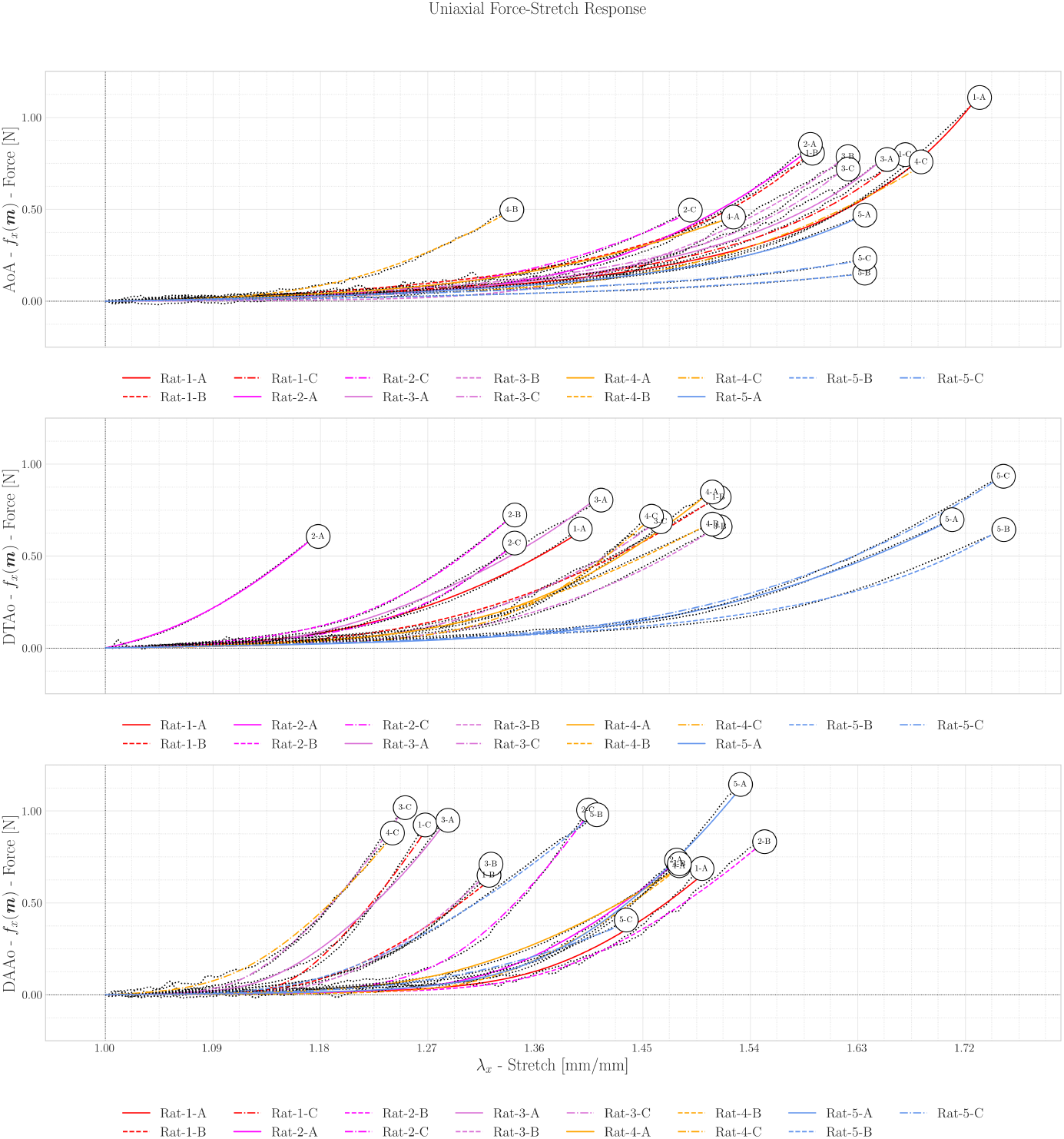
Uniaxial force-stretch responses across the aortic tree. Panels display the proximal ascending/aortic-arch (AOA), descending thoracic (DTAo), and abdominal (DAAo) segments for five subjects. Solid lines represent the optimized HGO-C constitutive model predictions, overlaid against the discrete experimental observations (markers). Colors denote individual animal specimens.

As shown in Figure 4, the rings from the proximal ascending/aortic-arch segment (AOA) demonstrated a more compliant initial response, consistent with the measured proximal protein composition and with a relatively larger matrix-associated contribution at low stretch. A modest increase in force was typically observed for stretch ratios starting from *λ*_*x*_ = 1.0 to approximately *λ*_*x*_ = 1.3 − 1.4. With increasing stretch, the curves stiffened progressively, indicating earlier fiber recruitment in the distal aorta than in the proximal segment [56, 59]. This more compliant proximal response is consistent with the buffering role often associated with the Windkessel effect [61, 59].

In comparison, rings from the DTAo (Figure 4, middle panel) typically began to exhibit significant stiffening at slightly lower stretch ratios. The transition to collagen-dominated behavior often appeared to initiate around *λ*_*x*_ = 1.2 − 1.3, with a subsequent sharp increase in the force. This suggests an earlier engagement of collagen fibers in the thoracic region than in the more proximal segments, indicating a shift in the microstructural composition or fiber architecture toward a less extensible tissue. This reduced compliance in the thoracic aorta reflects its evolving role from primarily a buffering vessel to more of a conduit, requiring increased stiffness to withstand propagating pressure waves [56].

The mechanical behavior of the rings from the infrarenal DAAo (Figure 4, bottom panel) generally showed the earliest onset of stiffening and the highest overall stiffness of the three regions studied, with force often increasing at stretch ratios as low as *λ*_*x*_ = 1.15 − 1.25. The increased stiffness and earlier recruitment of collagen in the DAAo align with the composition of a more muscular wall and smaller diameters (Table 1) and its physiological function in regulating peripheral resistance and distributing blood flow under higher mean and pulse pressures, where structural integrity and resistance to overstretching are paramount [32]. Considerable inter-sample variability was observed in this region. Overall, these regional differences in force-stretch behavior are consistent with the known proximal-to-distal stiffening gradient and spatial heterogeneity between different wall layers and along the length of the aorta [2, 56, 46].

### 4.3 Stress-Strain Analysis

To elucidate the mechanical behavior of the aortic wall, its stress contributions are decomposed into isotropic (matrix) and anisotropic (fiber) components based on the first Piola–Kirchhoff (PK1) stress tensor. This decomposition is not arbitrary; it derives directly from the fitted parameters of the implemented compressible Holzapfel–Gasser–Ogden (HGO-C) constitutive model.

Figure 5 presents the total PK1 stress (*P*_x_ = *P*_x-iso_ + *P*_x-fib_), alongside its constituent parts, for the proximal ascending/aortic-arch segment (AOA). Figures 6 and 7 detail the descending thoracic (DTAo) and abdominal (DAAo) segments, respectively. Baseline parameter responses appear in Figure 8. Across regions, the stress decomposition shows a qualitatively similar transition from a more matrix-associated low-stretch response to a more fiber-dominated high-stretch response, although the magnitude and onset of stiffening vary across animals and segments.

**Figure 5:**
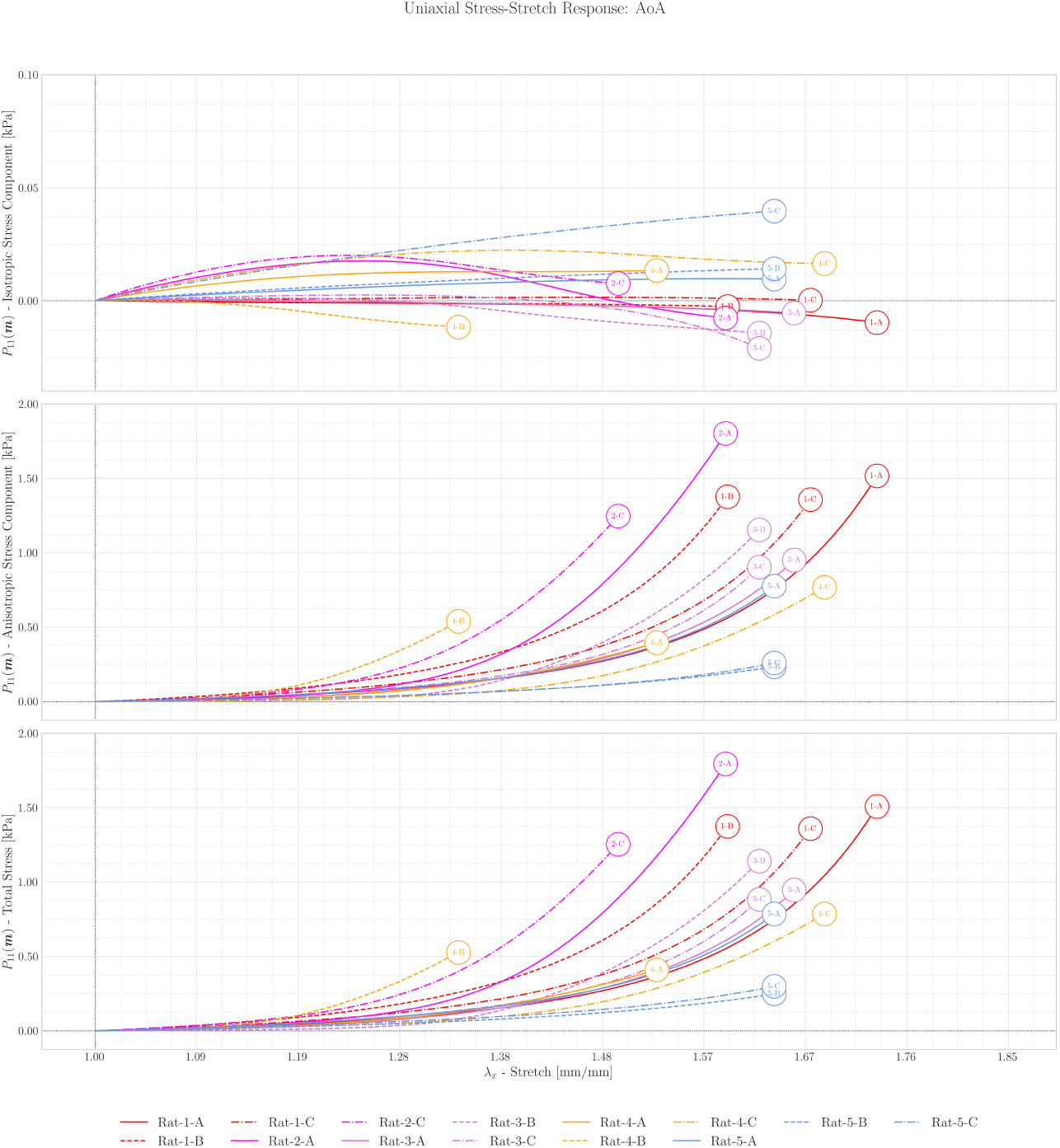
Decomposed stress-stretch behavior of the proximal ascending/aortic-arch (AOA). The single-layer HGO-C model resolves the total stress into its isotropic and anisotropic constituents using locally refined parameters. Distinct colors indicate different rat specimens (refer to Figure 4 for ID).

**Figure 6:**
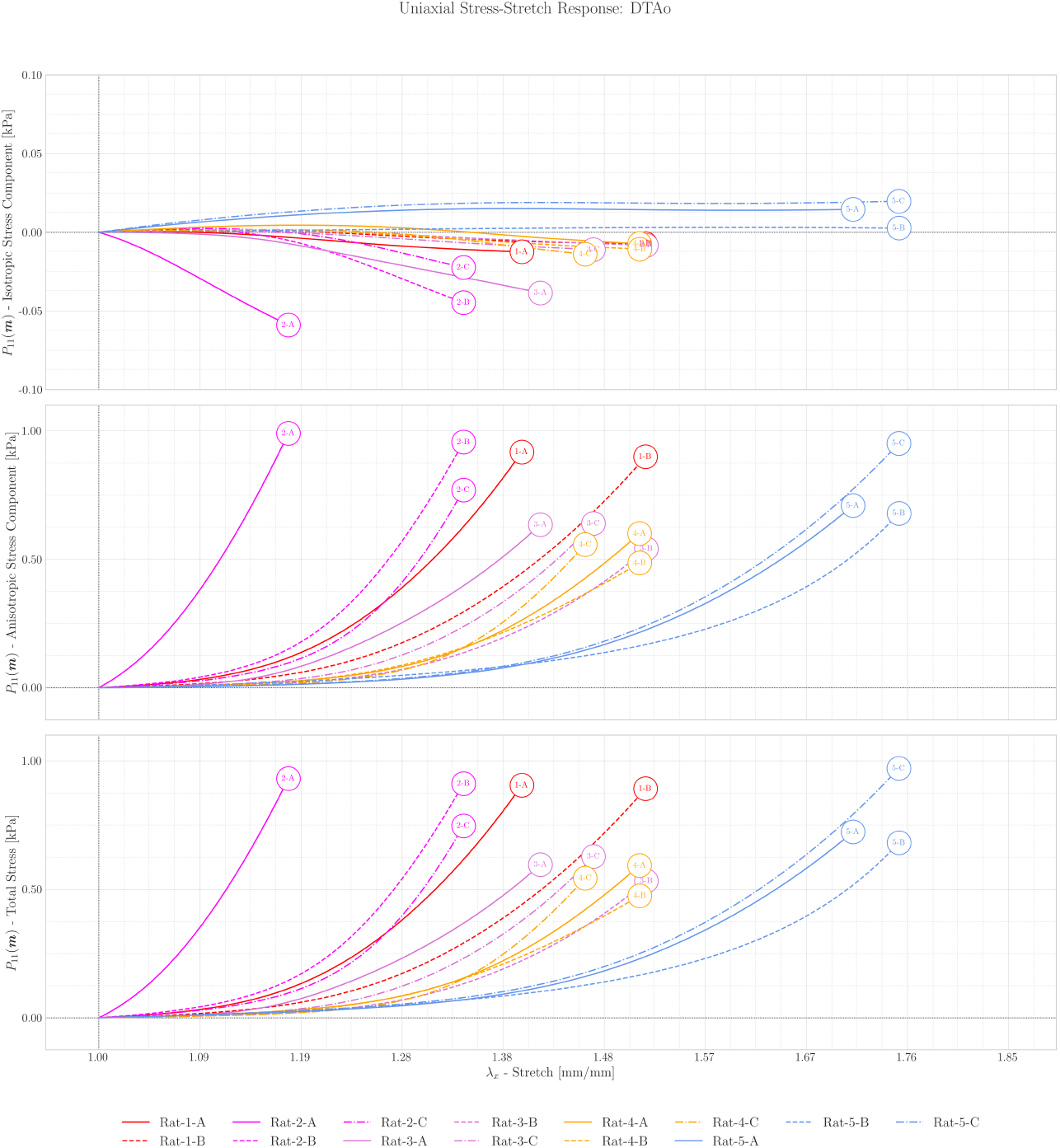
Stress-stretch curves (isotropic, anisotropic, and total components) of the single-layer model for Descending Thoracic Aorta (DTAo) ring segments from different rats, based on locally refined parameters.

**Figure 7:**
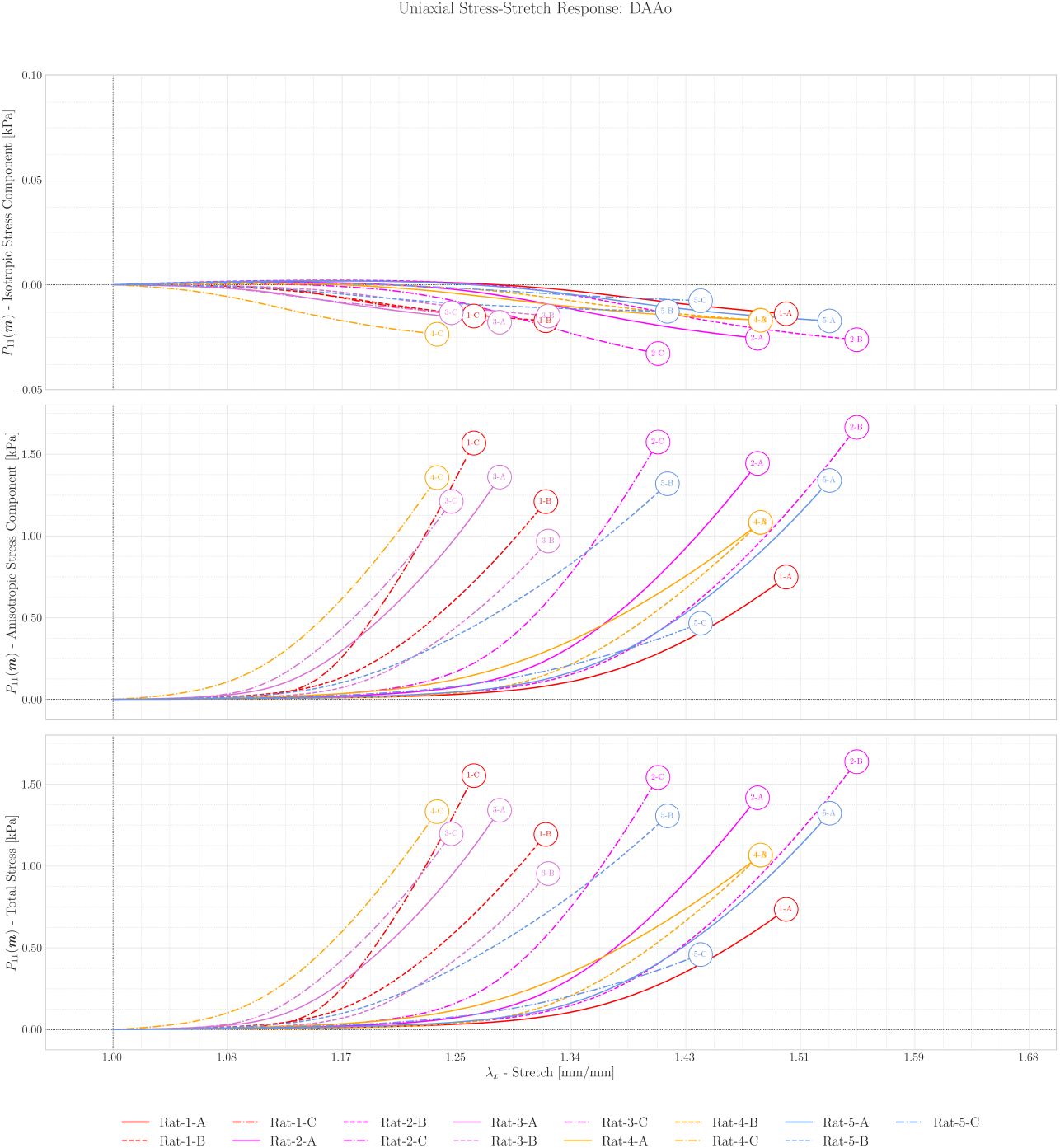
Stress-stretch curves (isotropic, anisotropic, and total components) of the single-layer model for Descending Abdominal Aorta (DAAo) ring segments from different rats, based on locally refined parameters.

**Figure 8:**
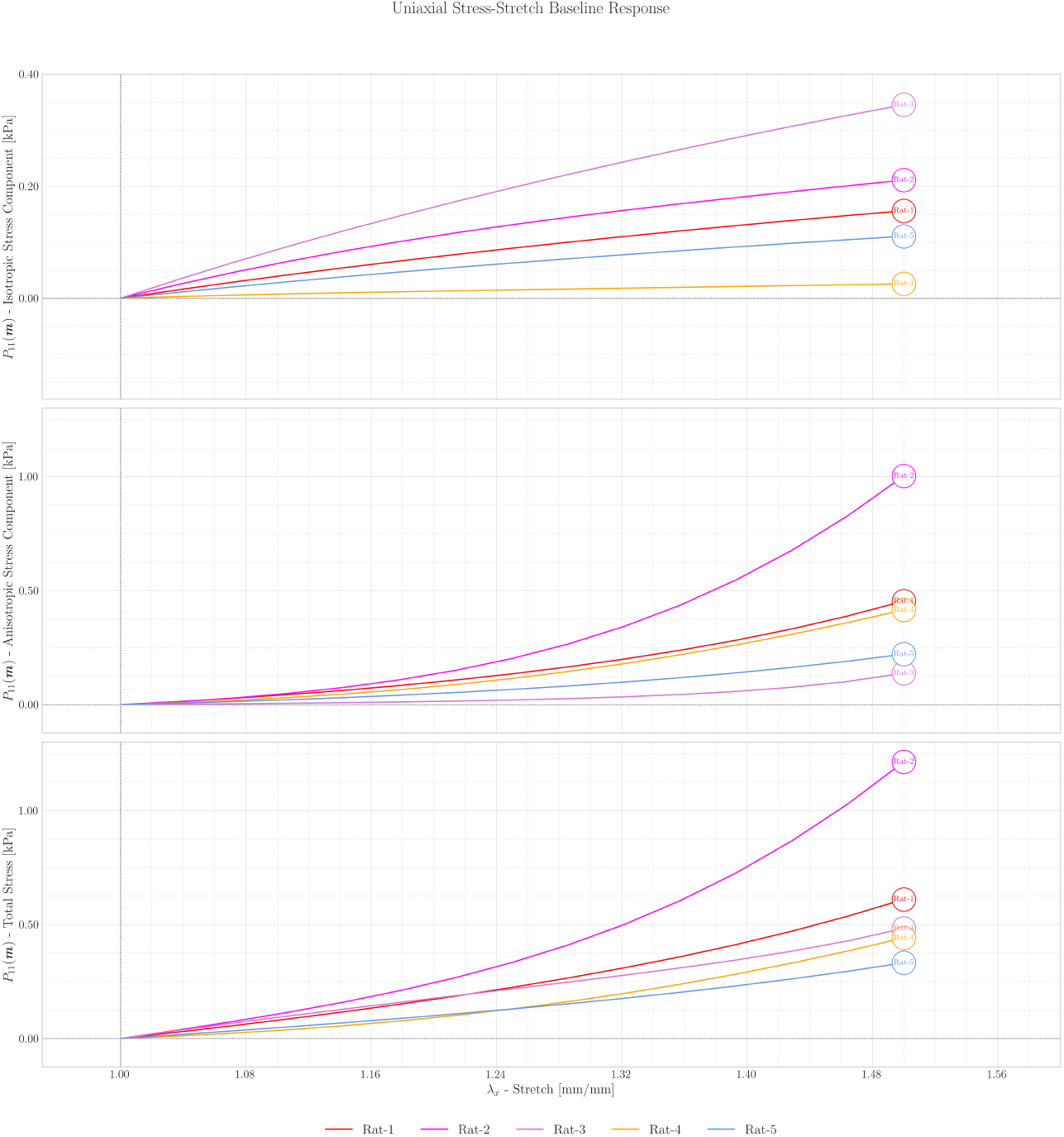
Uniaxial Stress-Stretch Baseline Response: Isotropic, Anisotropic, and Total stress components predicted using the baseline material parameters for each of the 5 rats

#### Isotropic Contribution (*P*_x-iso_)

The isotropic component, interpreted here as the model’s matrix-associated contribution, remains small relative to the total stress but varies across regions and stretch levels. In the proximal ascending/aortic-arch segment (AOA), this term is more variable at low stretch, whereas distal segments remain near zero or negative under the present uniaxial-fitting assumptions. These trends are consistent with regional compliance differences, but they should not be interpreted as direct evidence of a discrete elastin-to-collagen transition or a unique underlying microstructural mechanism.

When *P*_x-iso_ becomes negative, interpret the fitted decomposition with caution. In this constitutive model, these negative values show the balance between matrix-associated and fiber-associated terms under uniaxial loading. These values do not represent direct measurements of specific constituent-level events [22]. In the descending thoracic and abdominal regions, the consistently negative or near-zero isotropic contribution is therefore best interpreted as evidence of this modeled shift, not as proof of a unique physiological handoff mechanism [36].

#### Anisotropic (Fiber) Contribution (*P*_x-fib_)

The anisotropic contribution, represented by the fiber term in the constitutive model, dominates the mechanical response at higher stretch. In the fitted stress decomposition, this component’s magnitude is about two orders of magnitude larger than the matrix-associated one [34, 11]. As stretch increases, fiber recruitment rapidly raises stress. A clear regional trend emerges: high-stress transitions at lower stretch ratios in the DAAo than in the AOA; the DTAo shows an intermediate response. This behavior matches the stiffer distal aorta and its earlier fiber-associated response during deformation [59].

#### Total Stress (*P*_x_)

The total stress curves for all segments display the characteristic J-shaped behavior typical of soft biological tissues. The results show a proximal-to-distal stiffening gradient, with the AOA segments being the most compliant and the DAAo segments being the stiffest [56, 2]. The stress decomposition analysis suggests how this physiological gradient is represented within the fitted model. At low stretch, the proximal segment shows a relatively larger matrix-associated contribution and a more gradual stress rise, whereas with increasing stretch the response in all regions becomes increasingly fiber-dominated, with earlier recruitment in the distal aorta.

### 4.4 Material Parameter Estimation

Table 3 details the estimated material parameters ***m*** = {***m***_iso_, ***m***_fib_} = {*µ, D, β, κ, k*_1_, *k*_2_} for each aortic segment, derived via the dual-estimation strategy. The **baseline** parameters (one set per rat) capture the subject-specific characteristic elastic behavior, serving as a regularized prior for the subsequent local refinement. Consequently, the segment-specific parameters reflect regional mechanical variations and quantify the impact of microstructural heterogeneity along the vessel. In Table 3, *κ* [−] denotes the fiber dispersion factor (Equation (8)), ranging from perfectly aligned (*κ* = 0) to isotropic (*κ* = 1*/*3); for clarity, this is also expressed as a percentage of the isotropic limit (*κ*_max_ = 1*/*3): *κ* [%] = 100 *κ/κ*_max_. Where the parameter *k*_2_ approached zero (observed in specific baseline and local estimates), the exponential term in Equation (13) was evaluated via the limit *k*_2_ → 0 simplifying the fiber strain energy to a quadratic form 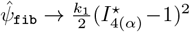, or simply 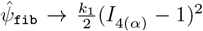 when *κ* = 0. Figure 9 illustrates the longitudinal trends of these refined parameters, using boxplots to summarize inter-animal variability and a grey line to highlight the mean trend across the cohort. Finally, the stress-strain responses corresponding to the baseline parameters for each rat are presented in Figure 8.

**Table 3:** Estimated material parameters in AOA, DTAo and DAAo segments in the studied animals.

| Rat N. | Section | Segment | $\mu$ [kPaPa] | $D$ [kPaPa] | $k_1$ [kPaPa] | $k_2$ [-] | $\beta$ [rad] | $\beta$ [deg] | $\kappa$ [-] | $\kappa$ [%] |
| --- | --- | --- | --- | --- | --- | --- | --- | --- | --- | --- |
| 1 | baseline | - | 0.1833 | 0.3855 | 0.1549 | 0.4983 | 0.5983 | 34.2801 | 0.0484 | 14.5127 |
| 1 | AOA | A | 0.0021 | 0.3974 | 0.0399 | 0.5983 | 0.3966 | 22.7235 | 0.0086 | 2.5905 |
| 1 | AOA | B | 0.0029 | 0.3905 | 0.0703 | 0.6700 | 0.3532 | 20.2369 | 0.0000 | 0.0000 |
| 1 | AOA | C | 0.0065 | 0.3926 | 0.0509 | 0.6115 | 0.3901 | 22.3511 | 0.0000 | 0.0000 |
| 1 | DTAo | A | 0.0210 | 0.4306 | 0.7480 | 0.6935 | 0.6368 | 36.4860 | 0.0000 | 0.0000 |
| 1 | DTAo | B | 0.0196 | 0.4372 | 0.5790 | 0.5002 | 0.6962 | 39.8893 | 0.0000 | 0.0000 |
| 1 | DTAo | C | - | - | - | - | - | - | - | - |
| 1 | DAAo | A | 0.0098 | 0.3725 | 1.2967 | 0.6358 | 0.7773 | 44.5360 | 0.0002 | 0.0599 |
| 1 | DAAo | B | 0.0049 | 0.4008 | 1.2924 | 0.7384 | 0.5811 | 33.2946 | 0.0000 | 0.0000 |
| 1 | DAAo | C | 0.0015 | 0.3924 | 2.1112 | 0.8110 | 0.5420 | 31.0543 | 0.0002 | 0.0536 |
| 2 | baseline | - | 0.3022 | 1.1241 | 1.7724 | 0.2809 | 0.8300 | 47.5555 | 0.0000 | 0.0000 |
| 2 | AOA | A | 0.0582 | 1.1139 | 2.3365 | 0.5168 | 0.8160 | 46.7534 | 0.0000 | 0.0000 |
| 2 | AOA | B | - | - | - | - | - | - | - | - |
| 2 | AOA | C | 0.0803 | 1.1309 | 1.9588 | 0.3264 | 0.7817 | 44.7881 | 0.0000 | 0.0000 |
| 2 | DTAo | A | 0.0826 | 1.1612 | 1.7554 | 0.2588 | 0.4781 | 27.3931 | 0.0619 | 18.5628 |
| 2 | DTAo | B | 0.0470 | 1.1470 | 1.7819 | 0.2700 | 0.6979 | 39.9867 | 0.0272 | 8.1460 |
| 2 | DTAo | C | 0.0352 | 1.1288 | 1.8983 | 0.2903 | 0.7079 | 40.5597 | 0.0032 | 0.9649 |
| 2 | DAAo | A | 0.0113 | 1.1281 | 1.9347 | 0.3317 | 0.7394 | 42.3645 | 0.0000 | 0.0000 |
| 2 | DAAo | B | 0.0124 | 1.1284 | 2.0067 | 0.3217 | 0.7772 | 44.5303 | 0.0001 | 0.0388 |
| 2 | DAAo | C | 0.0114 | 1.1193 | 2.5923 | 0.4417 | 0.7039 | 40.3305 | 0.0006 | 0.1765 |
| 3 | baseline | - | 0.3456 | 0.9572 | 0.0100 | 1.3511 | 0.4782 | 27.3988 | 0.0000 | 0.0000 |
| 3 | AOA | A | 0.0086 | 0.9592 | 0.1451 | 1.3752 | 0.6599 | 37.8095 | 0.0000 | 0.0000 |
| 3 | AOA | B | 0.0069 | 0.9584 | 0.7381 | 1.4369 | 0.7717 | 44.2152 | 0.0002 | 0.0491 |
| 3 | AOA | C | 0.0186 | 0.9559 | 0.0669 | 1.3449 | 0.4438 | 25.4279 | 0.0990 | 29.7056 |
| 3 | DTAo | A | 0.0067 | 0.9669 | 0.4521 | 1.0465 | 0.6469 | 37.0646 | 0.0421 | 12.6230 |
| 3 | DTAo | B | 0.0094 | 0.9582 | 0.4410 | 1.3407 | 0.7325 | 41.9692 | 0.0000 | 0.0000 |
| 3 | DTAo | C | 0.0101 | 0.9580 | 0.5508 | 1.3456 | 0.7017 | 40.2044 | 0.0000 | 0.0001 |
| 3 | DAAo | A | 0.0033 | 0.9579 | 1.1870 | 1.4901 | 0.5220 | 29.9084 | 0.0005 | 0.1507 |
| 3 | DAAo | B | 0.0029 | 0.9569 | 1.1038 | 1.5449 | 0.5956 | 34.1254 | 0.0004 | 0.1089 |
| 3 | DAAo | C | 0.0018 | 0.9578 | 1.1410 | 1.4655 | 0.4819 | 27.6108 | 0.0004 | 0.1199 |
| 4 | baseline | - | 0.0517 | 0.6835 | 0.2438 | 0.0100 | 0.6775 | 38.8179 | 0.0220 | 6.5862 |
| 4 | AOA | A | 0.0378 | 0.6886 | 0.5238 | 0.0313 | 0.7953 | 45.5673 | 0.0000 | 0.0000 |
| 4 | AOA | B | 0.0112 | 0.6877 | 0.6473 | 0.0470 | 0.6254 | 35.8328 | 0.0000 | 0.0000 |
| 4 | AOA | C | 0.0437 | 0.6648 | 1.1919 | 0.1723 | 0.8789 | 50.3573 | 0.0000 | 0.0000 |
| 4 | DTAo | A | 0.0226 | 0.6882 | 0.8233 | 0.0185 | 0.7757 | 44.4443 | 0.0024 | 0.7134 |
| 4 | DTAo | B | 0.0133 | 0.6949 | 0.4417 | 0.0100 | 0.7353 | 42.1296 | 0.0103 | 3.0928 |
| 4 | DTAo | C | 0.0117 | 0.6811 | 1.0908 | 0.0608 | 0.7649 | 43.8255 | 0.0019 | 0.5645 |
| 4 | DAAo | A | 0.0137 | 0.6905 | 1.0023 | 0.1962 | 0.7069 | 40.5024 | 0.0000 | 0.0000 |
| 4 | DAAo | B | 0.0049 | 0.6814 | 1.3977 | 0.1996 | 0.7345 | 42.0838 | 0.0002 | 0.0555 |
| 4 | DAAo | C | 0.0112 | 0.6778 | 1.5615 | 0.2552 | 0.4949 | 28.3557 | 0.0000 | 0.0000 |
| 5 | baseline | - | 0.1284 | 0.2349 | 0.1111 | 0.5371 | 0.6888 | 39.4653 | 0.0422 | 12.6670 |
| 5 | AOA | A | 0.0149 | 0.2499 | 0.0403 | 0.6715 | 0.4295 | 24.6085 | 0.0000 | 0.0000 |
| 5 | AOA | B | 0.0174 | 0.2430 | 0.0404 | 0.5718 | 0.6171 | 35.3572 | 0.0000 | 0.0001 |
| 5 | AOA | C | 0.0379 | 0.2454 | 0.0242 | 0.6157 | 0.5360 | 30.7105 | 0.0000 | 0.0000 |
| 5 | DTAo | A | 0.0327 | 0.3222 | 0.6632 | 0.6113 | 0.8517 | 48.7988 | 0.0000 | 0.0000 |
| 5 | DTAo | B | 0.0068 | 0.2524 | 0.0304 | 0.5907 | 0.5082 | 29.1177 | 0.0000 | 0.0000 |
| 5 | DTAo | C | 0.0389 | 0.3399 | 0.7647 | 0.6380 | 0.8572 | 49.1139 | 0.0000 | 0.0000 |
| 5 | DAAo | A | 0.0116 | 0.2767 | 1.5604 | 0.8967 | 0.7644 | 43.7969 | 0.0000 | 0.0000 |
| 5 | DAAo | B | 0.0038 | 0.3063 | 0.7228 | 0.3834 | 0.5724 | 32.7961 | 0.0000 | 0.0000 |
| 5 | DAAo | C | 0.0115 | 0.2912 | 0.5708 | 0.5930 | 0.7122 | 40.8061 | 0.0000 | 0.0000 |

**Figure 9:**
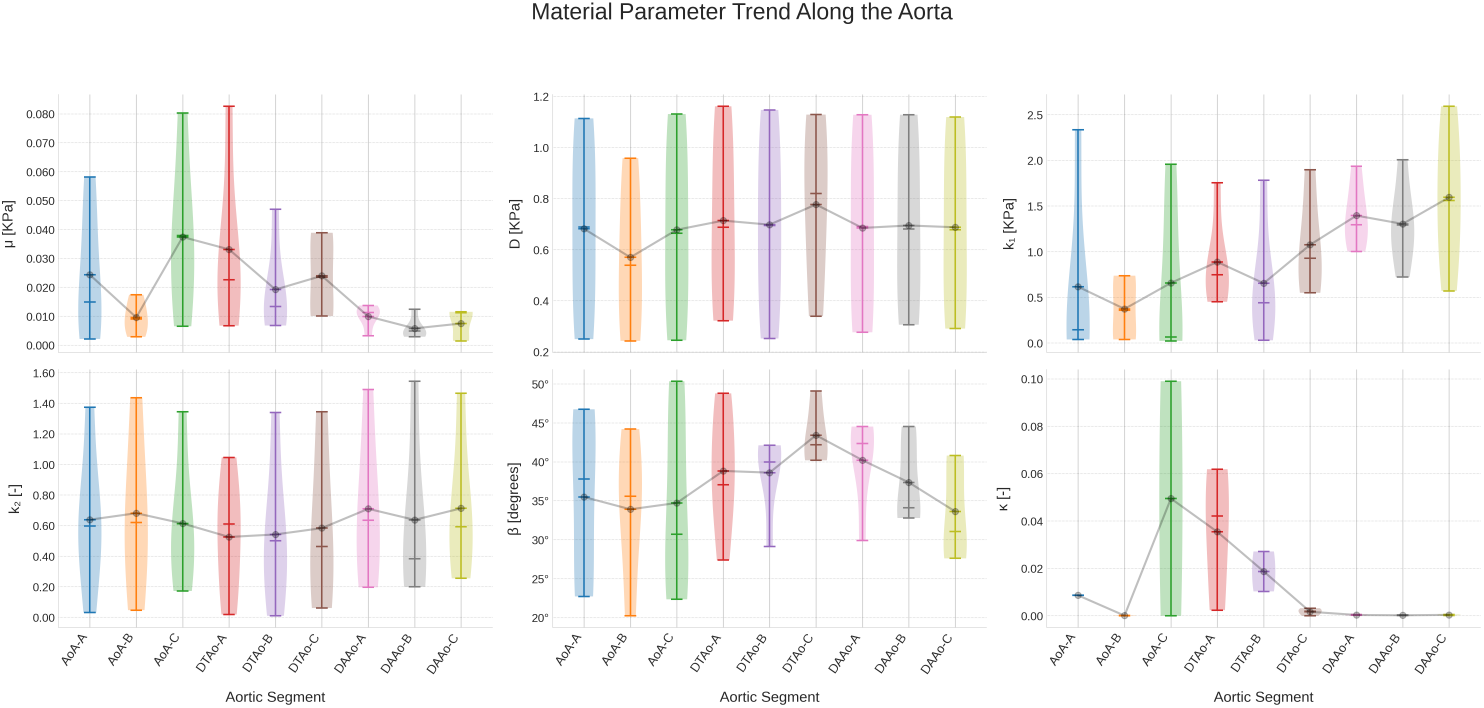
Regional distribution of locally refined constitutive parameters. The variation of *µ, D, k*_1_, *k*_2_, *β*, and *κ* is mapped across sequential aortic segments (e.g., AOA-A through DAAo-C). Box plots quantify the aggregate variance for the cohort (*N* = 5), while colored lines isolate specific subject trajectories. The grey overlay indicates the average parameter progression along the vessel length.

Several observations can be made regarding the estimated material parameters by analyzing the mean trends (grey lines in Figure 9):

- **Shear Modulus** (*µ*): The isotropic ground matrix parameter *µ* exhibits a distinct proximal-to-distal decay, reflecting a softer non-collagenous matrix in the abdominal regions. The mean trend is an initial increase within the arch (AOA-A to AOA-B/C) followed by a steady decline towards the distal DAAo segments. This gradient means that as one moves distally, the ground matrix offers less resistance to deformation. As a result, the load-bearing responsibility shifts almost entirely to the stiffening collagen fibers. This shift is consistent with the conduit function of the distal aorta. The lower initial value at AOA-A may reflect anatomical features of the aortic root or inter-animal variability. However, the overall trend remains robust. Elastin content remained regionally similar in our cohort. Thus, the variation in *µ* likely reflects differences in elastin organization, cross-linking quality, or the contribution of other matrix components like proteoglycans [56]. Although species differences complicate direct comparison, similar regional variations in matrix stiffness have been reported in human aortas [55].
- **Bulk Modulus** (*D*): The mean trend for *D* shows a tendency to decrease from proximal (AOA) to distal (DAAo) segments. As noted in Section 3.2, the fitted *D* values are regularization-influenced descriptors rather than direct estimates of the physical bulk modulus, because the volumetric regularization weight *E*_*b*_ largely governs quasi-incompressibility in our framework. Nevertheless, the proximal-to-distal trend suggests that the optimization allocates a marginally larger volumetric stiffness in the proximal aorta. However, the overall variability is considerable, and the physiological significance of this subtle mean trend needs further investigation, especially given the limitations of uniaxial testing in robustly determining *D*.
- **Collagen Parameters** (*k*_1_, *k*_2_): The mean trend for *k*_1_ (collagen stiffness-like parameter) exhibits a clear and substantial increase from the AOA through the DTAo to the DAAo, strongly indicating progressively stiffer collagen fibers or a greater effective contribution of collagen to tissue stiffness in the distal aorta, this aligns well with the physiological requirement for increased aortic stiffness distally [31] and the observed (though not statistically significant in our protein data) trend of increasing collagen I content [56]. The non-linearity parameter *k*_2_, which governs the rate of fiber uncrimping, peaks in the thoracic region, suggesting that the transition from the compliant “toe” region to the stiff linear region of the stress-strain curve is most abrupt in the thoracic aorta. For instance, Teng et al. [55] reported HGO parameters for human aorta layers, where *k*_1_ for adventitia was generally higher than for media, and *k*_2_ also varied.
- **Fiber Orientation Angle** (*β*): The mean trend for *β* demonstrates a more complex pattern than a simple monotonic increase. Specifically, the thoracic region (DTAo) exhibits larger fiber angles compared to both the ascending/arch (AOA) and abdominal (DAAo) regions. This elevated thoracic signal aligns with findings in larger mammals, where fiber architecture transitions along the aorta [56, 47]. Notably, this regional pattern is descriptive and pertains only to the non-calibrated subset of sections. Covariance analysis indicates that 17 of 43 sections require non-positive-definite Hessian calibration. These sections yield inconclusive region-level inference for *β*, and therefore, a cohort-wide monotonic proximal-to-distal trend cannot be established from uniaxial data alone.
- **Dispersion Parameter** (*κ*): The dispersion parameter *κ* remained consistently close to zero across all regions. This suggests highly aligned fibers. However, this interpretation requires caution. In a uniaxial tension test, fibers naturally reorient towards the loading axis. The optimization algorithm likely identifies this reoriented state as “aligned” (*κ* ≈ 0) to maximize the fiber contribution to the single available stress component (*P*_*xx*_). These results contrast with some studies on dissected murine aortas, in which regional *κ* values showed greater variation. Such variations could reflect different loading states or disease [5]. Thus, while *κ* accurately captures *effective* uniaxial behavior, it mainly reflects the recruited mechanical state. The actual morphological dispersion in the unloaded state is likely higher. Therefore, within this framework, *κ* should be interpreted as an effective descriptor of the loaded state. It should not be seen as a direct estimate of unloaded histological dispersion. As a result, parameter sets found this way should not be directly transferred to biaxial, inflation, or three-dimensional simulations without re-identification or independent structural constraints.

The dual-estimation strategy appears to yield parameter sets that produce relatively smooth variations along the aorta **within** each animal, as suggested by the individual animal trajectories in Figure 9. The standard deviations (indicated by the spread of boxplots) for many parameters, while present, are often comparable to or sometimes lower than those reported in literature for similar tissues characterized by HGO-type models from uniaxial or biaxial tests (e.g., SDs reported in [55] for human aorta or [1] for porcine aorta), suggesting that the proposed methodology can maintain a degree of consistency and avoid excessive parameter fluctuation while capturing regional differences.

## 5 Conclusions

In this study, we introduced and validated a novel methodology for assessing the constitutive material parameters of the aortic wall, employing a dual-estimation strategy. This strategy integrates a physically consistent compressible Holzapfel-Gasser-Ogden (HGO) model, informed by Modified Anisotropic (MA) principles to ensure appropriate coupling between volumetric and anisotropic responses [40]. This comprehensive approach facilitated a detailed characterization of the anisotropic invariants essential for accurately describing the aortic wall’s complex mechanical response under tensile loading [41, 18]. By systematically accounting for distinct material properties across different aortic regions ascending/arch, thoracic, and abdominal our methodology successfully enhanced the physiological fidelity of the characterized elastic behavior. The dual-estimation strategy, which first established animal-specific baseline parameters and subsequently refined these for individual aortic segments with regularization towards this baseline, yielded material parameters that, as observed in Figure 9 for individual animals, generally exhibited physiologically plausible and relatively smooth variations along the aortic length. This outcome aligns with a key objective of representing the aorta as a functionally integrated organ with graded, rather than abruptly changing, properties.

The two steps of global optimization, using the BH algorithm, effectively provide the baseline material parameters that capture the overall elastic signature of each rat’s aorta, allowing a second local optimization, which is used to refine these parameters for each specific aortic segment, leveraging analytical gradients computed via the adjoint method. The use of the L-BFGS algorithm within IPOPT contributed to computational efficiency. This combined global-local optimization approach demonstrated consistent robustness in navigating the complex objective functions characteristic of these inverse problems and supported stable estimation of the material parameters.

The main finding is that the estimated material parameters and decomposed stress contributions (isotropic *P*_x-iso_ and anisotropic *P*_x-fib_) closely matched regional differences in measured aortic responses. This reflects known variations in microstructure and fiber orientation. For example, the isotropic stress component *P*_x-iso_ in the ascending aorta usually stayed positive at initial stretches. It then dropped or turned negative at higher deformations, consistent with its compliant, elastin-driven role in the Windkessel effect [61]. In contrast, in stiffer descending thoracic and abdominal aortas, *P*_x-iso_ was often negative or near-zero, showing immediate reliance on the anisotropic fiber network. The collagen stiffness parameter *k*_1_ tended to increase distally, indicating growing aortic stiffness [31, 56]. Parameters like *µ* (matrix stiffness) and *D* (bulk modulus) generally decreased distally. The fiber angle *β* showed an elevated thoracic signal rather than a monotone proximal-to-distal increase. Dispersion *κ* stayed consistently low, reflecting high fiber alignment. These regional differences, as captured by our method, are important for understanding aortic function, such as its role in buffering pulsatile flow [59]. Tikhonov regularization, especially the baseline-guided step, stabilized the inverse problem. However, the resulting smooth parameter changes reflect the method rather than the underlying biology. These estimates capture regional trends but do not prove that biological gradients are as smooth as suggested by regularization. Effective alignment parameters from uniaxial fits should not be interpreted as direct histological measures.

### 5.1 Linking Biology to Mechanics

This study compares fitted HGO-C parameter trends with segment-specific quantification of extracellular matrix proteins in the rat aorta. By evaluating material-parameter trends in Figure 9 alongside protein quantification in Table 2, we assess whether regional mechanical patterns reflect underlying biological differences. These comparisons indicate that regional mechanical behavior is generally consistent with segment-specific protein composition. However, uniaxial inverse fitting cannot directly reconstruct unloaded microstructural architecture or capture all tissue complexity.

#### Collagen I and Fiber Stiffness (*k*_1_)

A necessary finding is the consistent relationship between collagen I concentration and the fiber stiffness parameter *k*_1_, rather than *k*_1_ serving as a direct surrogate for collagen content. Protein quantification shows that collagen I content rises from the proximal to distal aorta (AOA *<* DAAo, *P <* 0.05). The fitted material-parameter model reflects this gradient, with *k*_1_ also increasing along the same anatomical path. There is a strong positive correlation between mean collagen I intensity and estimated *k*_1_ across regions. This supports *k*_1_ as a meaningful mechanical correlate of collagen-related stiffening rather than a direct surrogate for collagen content. Higher distal *k*_1_ values and collagen I levels in the abdominal aorta align with earlier increases in anisotropic stress seen in Figure 7, matching the stiffer function of this region.

#### The Elastin Behavior Paradox

The relationship between elastin content and the matrix parameters (*µ* and *D*) reveals a more complex interaction and non-intuitive behavior. While the mean elastin content remained relatively constant or increased slightly towards the distal aorta (Table 2), the shear modulus *µ* (representing the ground matrix stiffness) generally decreased distally. This apparent discrepancy highlights the “Elastin Paradox”, in which protein abundance may not directly dictate matrix stiffness. Instead, the mechanical phenotype appears driven by the **ratio** of collagen to elastin. In the ascending aorta, the lower Collagen I/Elastin ratio corresponds to a lower *k*_1_*/µ* ratio in our model. This balance allows the isotropic matrix properties to dominate the low-strain regime, facilitating the Windkessel effect. Contrarily, in the abdominal aorta, the high Collagen I/Elastin ratio overwhelms the matrix contribution, shifting the tissue behavior to a fiber-dominated, anisotropic response.

Furthermore, the bulk modulus *D* showed a slight distal decrease but remained relatively robust; however, as discussed in Section 3.2, the fitted *D* values are regularization-influenced descriptors, so this trend should be interpreted as a regularization-informed observation rather than a direct measurement of regional compressibility. The stability of elastin content across different regions may explain why the bulk modulus *D* does not fluctuate as drastically as the fiber parameters. These findings suggest that *µ* in the HGO-C model captures the *effective* deviatoric stiffness of the non-collagenous matrix, which may be influenced by cross-linking density and proteoglycan interactions. While elastin’s intact architecture and volumetric constraint capability underpin the physical basis for *D*, in our uniaxial framework *D* primarily acts as a stabilizing model parameter ensuring thermodynamic consistency (see Section 3.2).

#### The Role of Collagen III

Interestingly, our data show that Collagen Type III content increases distally, similar to Type I. Since Collagen III is often associated with tissue compliance and extensibility, its accumulation in the stiffest region of the aorta (DAAo) seems counter intuitive. However, this suggests that the total collagen load (Type I + III) or the overriding stiffness of Type I bundles governs the macroscopic response. The increased Collagen III in the abdominal aorta may play a critical role in structural integrity and rupture prevention under high pressure, rather than compliance, acting as a safety net for the stiffer, less extensible wall.

#### Summary of Structure-Function Coupling

The dual-estimation strategy captured region-dependent mechanical trends by using a regularized inverse formulation to stabilize local parameter estimation. Furthermore, Western Blot results agreed with the trends of the fitted parameters, supporting that the identified regional variations are biologically meaningful, especially for parameters closely tied to the observed uniaxial response, such as *k*_1_, which reflects fiber stiffness. The smooth changes in Figure 9 should be interpreted as regularized trends consistent with regional biology. However, this does not prove that the underlying biological transition is continuous in the same form. The microstructural interpretation of *κ* (fiber dispersion), and to a lesser extent *β* (fiber orientation angle), remains more limited under uniaxial loading.

### 5.2 Limitations and Future Work

While this study advances robust identification of regional aortic mechanics, some limitations have been carefully considered. However, certain limitations must be addressed in future studies.

#### Uniaxial Loading vs. *In Vivo* Complexity

The primary limitation is the use of uniaxial ring-extension data to fit a 2D anisotropic model. As described in Simple tension, the ring’s response is measured with the circumferential load as the *x*-axis. The deformation gradient is diagonal, ***F*** = diag(*λ*_*x*_, *λ*_*y*_, *λ*_*z*_), and the PK1 stress tensor is given by *P*_*xx*_, *P*_*yy*_, and *P*_*zz*_. Here, the material parameters describe how the ring behaves circumferentially in the uniaxial test. This view assumes plane stress (*P*_*yy*_ = 0, *P*_*zz*_ = 0), with *P*_*xx*_ carrying the load. However, uniaxial tests cannot fully capture the multiaxial stress state or account for interactions between circumferential and longitudinal loads, as in vivo. This study focused on comparing regional differences rather than fully identifying the constitutive model. Thus, the same bias from the loading mode affects all samples. The reported pattern of greater stiffening from proximal to distal is a comparative trend in this dataset. Future work, such as biaxial or inflation-extension tests, could refine key parameters, such as the fiber angle *β* and the volumetric-response parameter *D*.

#### Interpretation of Fiber Dispersion (*κ*)

Our optimization consistently yielded low dispersion values (*κ* ≈ 0), indicating highly aligned fibers in the aortic arch. However, this opposes histology, which shows a more dispersed fiber network in proximal regions. This likely occurs because uniaxial tests cause fibers to realign toward the load at high strains. Thus, the low *κ* values reflect this reoriented state, not the natural orientation. The optimization minimizes misfit in the single force component 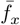 and so selects this test-induced alignment. Our modified compressible model removes the non-physical mathematical link between volume changes and fiber stretches seen in standard HGO-C models [40], but fitting *κ* to 1D macroscale data still presents phenomenological risk. To separate testing effects from true orientation, future work could take two approaches. Experimentally, shifting to multiaxial tests (e.g., biaxial or inflation-extension) would capture the 3D deformation field and prevent the algorithm from adjusting for unmeasured strains [39]. Microstructurally, *κ* could be directly bounded or fixed using optical measurements of fiber orientation (such as Second-Harmonic Generation microscopy or polarized light imaging), instead of macroscopic fitting alone.

#### Tikhonov regularization (ℛ_tik_)

The local Tikhonov term with *ω*_*r*_ = 10^−3^ should be interpreted as a guiding regularization term around the baseline solution rather than as a constraint that forces strict homogeneity. Although the fitted curves in Figure 4 remain segment-specific under this choice, any nonzero Tikhonov weight introduces some degree of shrinkage toward the reference solution. A formal sensitivity analysis with respect to *ω*_*r*_ was not performed in the present study and remains an important topic for future work.

#### The “Elastin Paradox” and Volumetric Constraints

Negative isotropic stress in the HGO-C model demonstrates the challenge of modeling fiber-filled materials with minimal compressibility using just uniaxial test data. The bulk modulus *D* here depends on movement data, which does not include information about side-to-side shrinkage. Without more limits, the HGO-C model allows a range of *D* values that satisfy the equations. We estimate *D* using a volume regularization term to ensure the results stay realistic and match those of materials with slight compressibility.

This modeling constraint approach highlights a critical link between bulk *D* and the shear modulus *µ*. This relationship helps clarify the “Elastin Paradox”, the gap between elastin’s biochemical abundance and its mechanical contribution. Biologically, the elastin-rich ground matrix serves two main functions: preserving volumetric constraint and acting as “glue” for collagen fibers. Our Western Blot data indicate constant elastin content along the vessel; however, regional variations in *µ* suggest that protein quantification alone does not strictly dictate tissue stiffness. Microstructural studies corroborate this perspective, showing that elastin’s functional stiffness depends on its intact architecture [19, 56]. Distal regions, such as the abdominal aorta, are more susceptible to elastin fragmentation, mechanical fatigue, and lamellar discontinuity [56, 23]. This decreases elastin’s mechanical competence as measured biochemically. Consequently, *D* is constrained to enforce physiological volume conservation, while regional differences in matrix integrity are mainly reflected in *µ*. Additionally, as the matrix degrades, the stiff collagen network engages at lower macroscopic stretches [23]. This behavior diminishes the isotropic matrixs relative contribution in 1D parameter estimation and forces a lower *µ* fit. Therefore, *D* acts as a stabilizing model parameter, whereas *µ* reflects the effective structural stiffness of the intact inter-fibrillar matrix, rather than simply the total elastin quantity.

#### Future Work

This work lays the groundwork for subject-specific modeling in vascular disease. Future work could test whether the present framework remains informative in pathological models, such as aneurysms or dissections, where regional property gradients may be altered.

### 5.3 Concluding Remarks

Overall, the dual-estimation strategy combines a compressible HGO formulation, modified-anisotropic (MA) ideas, and robust optimization into a practical framework for estimating aortic material parameters from uniaxial data while preserving subject-specific and regional trends. In that sense, the method offers regularized descriptors of regional mechanics rather than a complete physiological reconstruction.

This study helps clarify how regional mechanical trends in the aorta relate to measured protein composition and tissue behavior. Generating regional, smoothly varying material parameters can support comparative simulations of aortic hemodynamics and mechanobiology, but clinical inference would require additional validation and richer experimental loading data.

## Declaration of Generative AI in the Writing Process

During the preparation of this work, the authors used a generative AI (Grammarly and Claude) to improve the manuscript’s grammatical accuracy and readability. After using this tool, the authors reviewed and edited the content as needed and take full responsibility for the publication’s content.

## Funding

This work was supported by the Fundação de Amparo à Pesquisa do Estado de São Paulo (FAPESP) under grants 2012/50283-6, 2019/21236-9, 2023/03079-9, and 2025/27076-4.

The Article Processing Charge for the publication of this research was funded by the Coordenação de Aperfeiçoamento de Pessoal de Nível Superior - CAPES (ROR identifier: 00x0ma614). For open access purposes, the authors have applied a Creative Commons CC BY license to any accepted version of the article.

## Appendix A Experimental procedure

Aortic tissue samples were homogenized using a Precellys Evolution Touch (Bertin Technologies) and subsequently lysed in RIPA Lysis Buffer (Millipore), supplemented with protease and phosphatase inhibitors to preserve protein integrity. Following centrifugation at 12,000 xg for 15 minutes at 4°*C*, the supernatant containing the solubilized proteins was carefully collected. Aliquots of cell lysates (20 *µ*g total protein) were separated based on molecular weight using SDS-polyacrylamide gel electrophoresis (SDS-PAGE). The separated proteins were then electrotransferred to polyvinylidene difluoride (PVDF) membranes (Millipore). To prevent non-specific antibody binding, these membranes were incubated for 1 hour at room temperature in a blocking buffer solution, which consisted of 5% Bovine Serum Albumin (BSA), 10 mM Tris-HCl (pH 7.6), 150 mM sodium chloride, and 0.1% Tween 20. After blocking, the membranes were probed with specific primary antibodies: Collagen Type I (Abcam, ab34710) and Collagen Type III (Abcam, ab7778). Protein quantification in ascending, thoracic, and abdominal aortic segments was performed using this Western Blot analysis. The resulting protein bands were visualized using SuperSignal West Pico PLUS Chemiluminescence Substrate (Thermo Fisher Scientific) and subsequently quantified densitometrically using ImageJ software [43].

## Appendix B Derivation of the Cost Function Gradient

In this appendix, we outline the analytical derivation of the gradient of the cost function 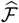 (23) with respect to the material parameters ***m***. This gradient is essential for the gradient-based optimization algorithms employed (Section 3.4). The core component involves the sensitivity of the predicted reaction force *f*_*x*_ to changes in the material parameters.

The gradient of the Cauchy loss term requires the partial derivative of the reaction force *f*_*x*_ with respect to the material parameters ***m***. Using the forward model definition (22) and assuming homogeneous deformation under simple tension where *P*_*xx*_ represents the relevant stress component and a^*r*^ is the reference area associated with one loading arm (total area 2a^*r*^), this derivative is

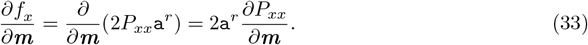

The derivative of the first Piola-Kirchhoff stress component, *P*_*xx*_ (which is derived from Eq. (21) using the constitutive model defined by Eqs. (5), (10), (11), and (13)), with respect to each material parameter in the set ***m*** = {*µ, D, k*_1_, *k*_2_, *β, κ*}, are given by the following expressions

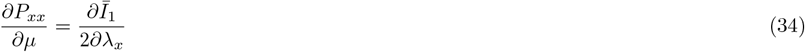

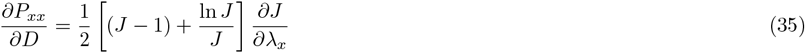

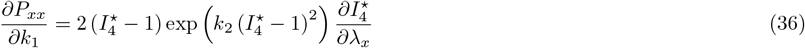

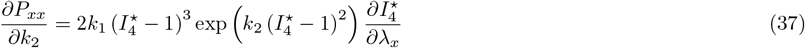

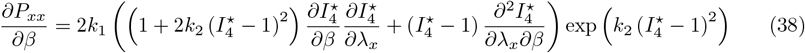

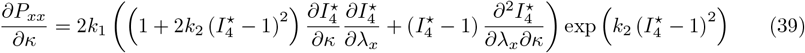

The gradient of the full cost function 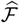 (23) is then computed by applying the chain rule to these sensitivities for the Cauchy loss term, and combining this with the direct derivatives of the regularization terms ℛ_tik_(***m***) and ℛ_vol_(***m***). The adjoint method, as outlined in Section 3.3, provides an efficient computational pathway to calculate the overall gradient required by the optimization algorithm.

## Appendix C Partial Derivatives of Strain Energy Components and Invariants

This appendix provides the necessary partial derivatives of the strain energy components (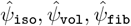) and the relevant kinematic invariants (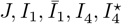) with respect to the principal stretches (*λ*_*i*_ for *i* ∈ {*x, y, z*}) or material parameters (*β, κ*). These derivatives are used in calculating the stress tensor (21) and the sensitivities required for the cost function gradient (Appendix B).

Derivatives of Strain Energy Components with respect to principal stretches

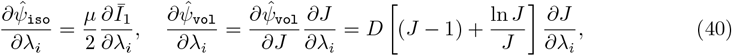

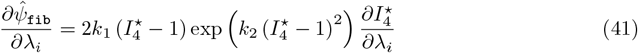

Derivatives of Invariants with respect to principal stretches, diag(***F***) = ***λ*** = [*λ*_*x*_, *λ*_*y*_, *λ*_*z*_]

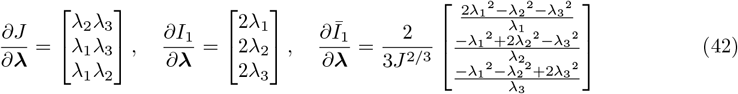

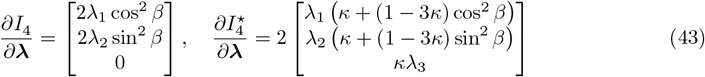

Derivatives of Fiber Invariants with respect to Material Parameters

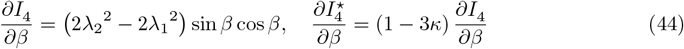

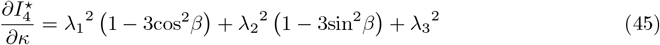

Second-Order Derivatives Needed for Sensitivity Calculations

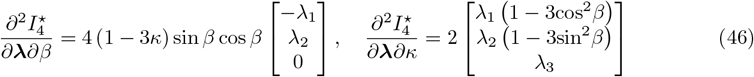

## Notes

https://github.com/Rlahuerta/dual-matfit.git

### Competing Interest Statement

The authors have declared no competing interest.

